# Single-cell transcriptomics of human astrovirus VA1-infected intestinal organoids reveals cell-type-specific responses and a requirement for cholesterol biosynthesis during infection

**DOI:** 10.64898/2026.11.17.767333

**Authors:** Andres Rivera Ruiz, Roberto J. Cieza, Ameer Almeshagbeh, Matthew McConnachie, Jonathan Z. Sexton, Elliot D. SoRelle, Christiane E. Wobus

## Abstract

Human astroviruses are prevalent but understudied enteric viruses that cause gastroenteritis and neurological disease, particularly in children and immunocompromised individuals. No vaccines or antivirals are available, partly because their biology and pathogenesis remain poorly understood. Here, we used single-cell RNA sequencing to characterize transcriptional responses to VA1, a non-classical human astrovirus associated with the majority of extraintestinal disease cases, in fetal ileum-derived human intestinal organoids. VA1 altered epithelial cell-type proportions and progressively spread through the organoids. Differential gene expression analysis of enterocytes and progenitor cells, the cells most prevalent in the organoid model, revealed shared antiviral responses, including interferon signaling, as well as cell-type-specific responses. Enterocytes exhibited a broader early transcriptional response and specifically upregulated cholesterol metabolism and biosynthesis pathways at 1 dpi. By 3 dpi, progenitor cell-specific genes suggested inflammatory and epithelial-defense responses. In two different intestinal epithelial cell lines, T84 and Caco-2 cells, limiting extracellular lipid availability did not alter replication but increased VA1 binding in Caco-2 cells. Pharmacologic inhibition of cholesterol biosynthesis and 25-hydroxycholesterol treatment reduced VA1 replication, while disrupting intracellular cholesterol transport had no detectable effect. Together, these findings expand our knowledge on HAstV biology by defining shared and cell-type-specific epithelial responses to VA1 infection and identifying cholesterol biosynthesis as an important host pathway for successful VA1 replication.

**Importance:** Human astroviruses are a common cause of gastroenteritis as well as severe neurological disease in some individuals, especially children and people with compromised immune systems. Despite astrovirus prevalence and clinical importance, no vaccines or targeted antiviral treatments are available, in part because key aspects of human astrovirus biology and pathogenesis remain poorly understood. How different cell types respond to infection currently constitutes a fundamental gap in knowledge. In this study we define shared and cell type-specific responses in human intestinal organoids to infection with human astrovirus VA1, the strain associated with most cases of encephalitis. We found that enterocytes and progenitor cells mount distinct responses to infection and identified cholesterol biosynthesis as a host process supporting efficient VA1 replication in enterocytes. These findings advance our understanding of human astrovirus infection and identify cholesterol biosynthesis as a potential target for developing antiviral strategies.

## Introduction

Human astroviruses (HAstVs) are positive-sense single-stranded RNA viruses that cause gastroenteritis, mostly in children, the elderly and immunocompromised people. HAstVs account for 2 to 9% of all acute non-bacterial gastroenteritis cases in children worldwide and for ∼24,600 deaths in 2023, mainly in children younger than 5 years of age (1–4). HAstVs are divided into two groups: classical and non-classical strains. Classical HAstVs, which include HAstV serotypes 1-8, have been conclusively linked to diarrhea (5). Their seroprevalence is high but variable between serotypes. HAstV-1 is the most prevalent serotype with antibody presence ranging from 73% to 100% among adults while pediatric seropositivity ranges from 94% to 100% in children aged 3 to 9 years (6–11). On the other hand, non-classical HAstVs (VA/HMO and MLB clades) have not been conclusively linked with gastrointestinal symptoms, but instead to neurological diseases, such as encephalitis (12). These cases tend to occur in immunocompromised patients and are often fatal (13). HAstV VA1 has been linked most frequently to HAstV-induced neurological disease (13). Non-classical HAstVs are also highly prevalent, with studies showing a seroprevalence of over 75% to VA1 in the United States and Peru, and a seroprevalence close to 100% for MLB1 in a United States cohort study (12, 14). Despite their high prevalence and public health burden, no antiviral treatments and/or vaccines are available against HAstVs due in part to the current knowledge gaps in general HAstV biology and pathogenesis.

The positive-sense RNA genome of HAstVs has an approximate length of 6.2 to 7.7 kilobases and contains three major open reading frames: ORF1a, ORF1b and ORF2. ORF1a encodes nonstructural proteins involved in remodeling of ER membranes into replication organelles for viral RNA replication, innate immune evasion and the viral protease (15–19). ORF1b encodes the RNA-dependent RNA polymerase, and ORF2 encodes the capsid protein (20). A fourth open reading frame, ORFX, which encodes a protein with viroporin function, is only found in classical and MLB strains (21). Following entry and uncoating, the genomic RNA is translated to produce two different polyproteins as the result of a ribosomal frameshift: Nsp1a, translated from ORF1a and Nsp1ab, translated from ORF1a and ORF1b, respectively. A negative-sense RNA intermediate is then generated and used as a template for the synthesis of new full-length genomic RNA. Early termination of the negative sense RNA generates the template for a shorter, highly abundant subgenomic RNA from which ORF2 is translated (22). Capsid proteins then assemble with genomic RNA into immature virions that subsequently mature before being released from the cell through cell lysis, extracellular vesicles, or mucus secretion from goblet cells (20, 23, 24).

In recent years, human intestinal organoids (HIOs) have become a popular model to study enteric virus infections, including HAstVs (25–27). HIOs are biopsy-derived cultures of intestinal epithelial cells that self-organize into three-dimensional structures. HIOs are non-transformed, contain multiple cell types (e.g., progenitor cells, enterocytes, goblet cells and enteroendocrine cells), and recapitulate characteristics of the intestinal segments from which they were derived. Investigations of HIOs have revealed HAstV-1 and VA1 can infect progenitor cells, goblet cells, and enterocytes (28, 29). However, cell-type-specific responses to HAstVs have only been investigated for classical HAstV-1 (29). Single-cell analysis of HAstV-1-infected ileum-derived intestinal organoids showed that enterocytes, stem/progenitor cells, transit-amplifying cells, and goblet cells mounted distinct antiviral transcriptional responses characterized by lineage-specific patterns of interferon (IFN) stimulated gene (ISG) expression. Basal ISG expression also differed among uninfected cell types, suggesting that each intestinal epithelial lineage exhibits a distinct intrinsic antiviral state and contributes differently to virus infection (29). Bulk transcriptomic analysis of VA1-infected HIOs showed strong type I and III IFN-driven antiviral responses with similarities and some differences to two other enteric viruses, namely human norovirus and rotavirus (30). However, single-cell transcriptomic analysis of the cell-type specific responses during VA1 infection, to our knowledge, has not been reported.

Therefore, we used single-cell RNA sequencing to characterize time- and cell-type-specific responses to VA1 infection in a fetal ileum-derived HIO line selected for its robust susceptibility to VA1 and donor age, as clinical symptoms predominantly occur in children (28, 31). VA1 transcripts were detected in all identified major cell types; enterocytes, progenitor cells, goblet cells and enteroendocrine cells. Shared and cell-type-specific responses to VA1 infection were compared between the two predominant cell types, enterocytes and progenitor cells. VA1 infection induced an antiviral innate immune response in both enterocytes and progenitor cells at 1 and 3 days postinfection (dpi). Enterocytes upregulated a broader set of genes involved in antiviral immunity than their progenitor counterparts at 1 dpi, while progenitor cells exhibited a gene signature suggestive of inflammatory and epithelial-defense responses by 3 dpi. Beyond immune genes, shared metabolic changes in both cell types at 3 dpi were suggestive of compromised digestive, detoxification, and regenerative capacities, while enterocytes, but not progenitor cells, upregulated genes involved in cholesterol homeostasis at 1 dpi. Follow-up studies in intestinal epithelial cell lines validated the role of cholesterol homeostasis during VA1 infection. Limiting extracellular lipid availability increased VA1 binding in Caco-2, but not T84, cells, while viral replication was unaffected in both cell lines, and no impact was observed when intracellular cholesterol transport was inhibited. On the other hand, pharmacological inhibition of cholesterol synthesis resulted in statistically significant decreases in viral titers in both cell lines. In addition, treatment with 25-hydroxycholesterol, an important negative regulator of cholesterol homeostasis, reduced VA1 replication. These data suggest the importance of cholesterol synthesis for optimal VA1 infection. Collectively, these findings describe the first analysis of cell type-specific responses to a non-classical HAstV strain and add offer new insights into the biology of these understudied, yet highly prevalent viruses.

## Results

### Unbiased clustering and annotation of single-cell fetal ileum-derived organoid data reveals cell types present in the intestinal epithelium

To study the response to VA1 infection at the cellular level, we inoculated fetal ileum-derived organoids with VA1 or Caco-2 mock lysates in a three-dimensional (3D) format, as previously described for human norovirus infections (32). Single-cell suspensions were collected for single-cell RNA sequencing at 1 and 3 dpi for each condition. After quality control processing and filtering, the mock dataset was used for unbiased clustering to avoid any influence of VA1-induced host responses on clustering. This analysis identified nine different cell populations: two enterocyte populations (clusters 1, 6), five progenitor-like populations (clusters 0, 2, 3, 5, 7), one goblet-cell population (cluster 4), and one enteroendocrine-cell population (cluster 8) **(Fig. S1A)**. To simplify downstream analyses, we grouped the multiple enterocyte- and progenitor-like populations into one enterocyte population and one progenitor cell population, respectively, for a total of four final cell populations **(Fig. S1B)**. We further confirmed these annotations by analyzing the expression of previously published cell type markers from a fetal small intestine epithelium single-cell dataset (33), identifying the main markers of each cell type cluster **(Table S1)**, and assessing their cell type-specific enrichment using the Human Protein Atlas dataset. **Fig. 1A** shows the relative expression of some of these markers identified for each annotated cell type cluster.

**Figure 1.**
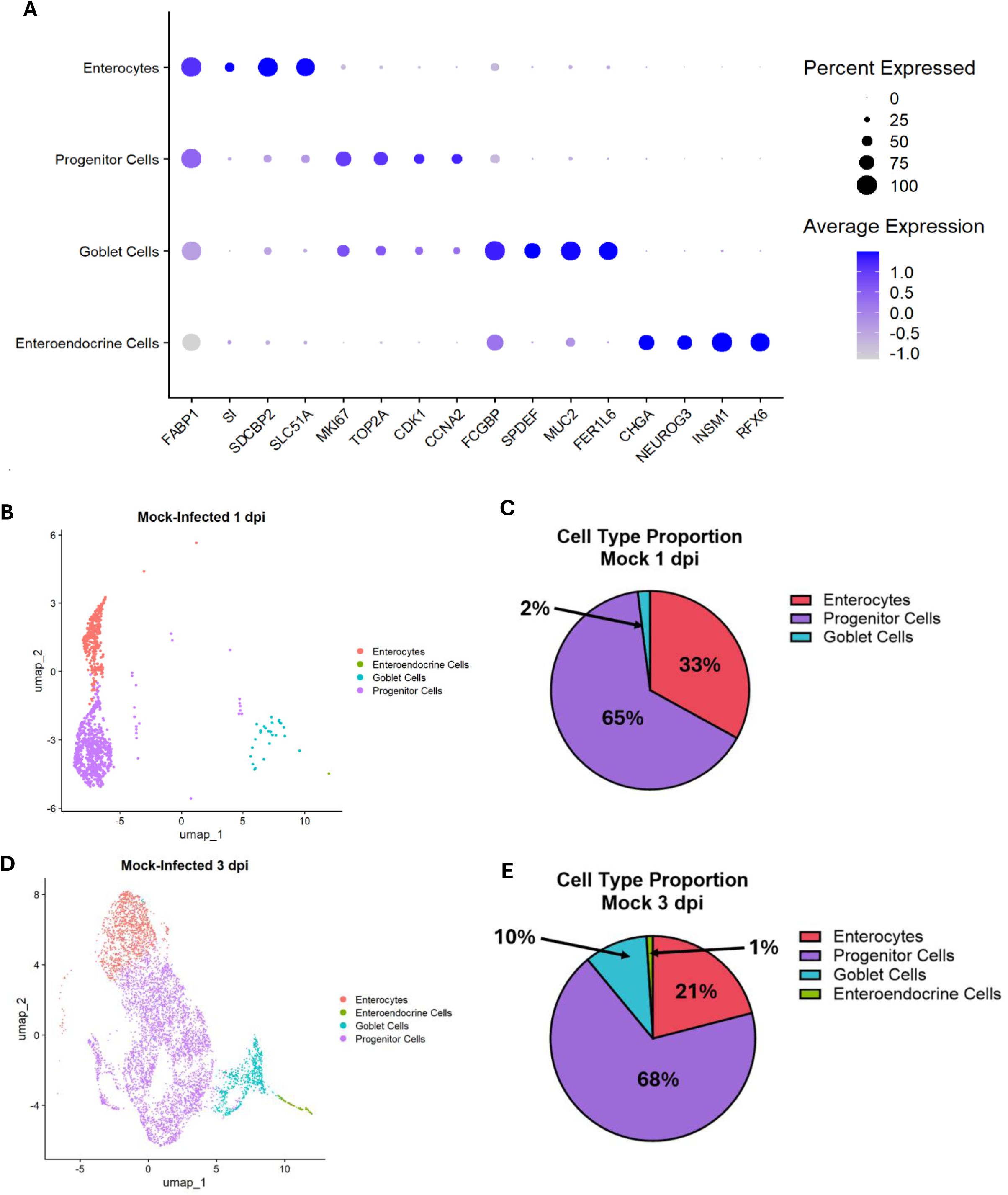
Unbiased clustering and annotation of single-cell fetal ileum-derived organoid reveals cell types present in the intestinal epithelium. **(A)** Dot plot showing the average expression of marker genes and the percentage of cells expressing each gene across the annotated cell-type clusters. Markers included fatty acid-binding protein 1 (*FABP1*), sucrase-isomaltase (*SI*), syndecan-binding protein 2 (*SDCBP2*), and solute carrier family 51 alpha subunit (*SLC51A*) for enterocytes; marker of proliferation Ki-67 (*MKIc7*), DNA topoisomerase II alpha (*TOP2A*), cyclin-dependent kinase 1 (*CDK1*), and cyclin A2 (*CCNA2*) for progenitor cells; Fc gamma-binding protein (*FCGBP*), SAM pointed domain-containing ETS transcription factor (*SPDEF*), mucin 2 (*MUC2*), and fer-1-like family member 6 (*FER1Lc*) for goblet cells; and chromogranin A (*CHGA*), neurogenin 3 (*NEUROG3*), INSM transcriptional repressor 1 (*INSM1*), and regulatory factor X6 (*RFXc*) for enteroendocrine cells. **(B, D)** Uniform manifold approximation and projection (UMAP) plots of mock-infected cells at 1 and 3 days postinfection (dpi), respectively, annotated by cell type. **(C, E)** Pie charts showing the distribution of cell types in mock-infected samples at 1 and 3 dpi, respectively.

After annotation, we assessed how epithelial cell type proportions changed over time in the mock condition. At 1 dpi, mock-infected organoids consisted of 65% progenitor cells, 33% enterocytes, and 2% goblet cells **(Fig. 1B-C)**. At 3 dpi, mock-infected organoids consisted of 68% progenitor cells, 21% enterocytes, 10% goblet cells, and 1% enteroendocrine cells **(Fig. 1D-E)**. The increased numbers of goblet and enteroendocrine cells at 3 dpi, suggests an increase in differentiated cells of intestinal epithelial organoids over time in culture.

Taken together, our analysis identified four major epithelial cell types in fetal ileum-derived organoids, with progenitor cells and enterocytes comprising most cells at both time points, while goblet and enteroendocrine cells expanded by 3 dpi.

### VA1 changes epithelial cell type proportions in infected human intestinal organoids

To determine whether VA1 infection altered epithelial cell-type proportions, we used Seurat’s anchor-based label-transfer workflow (34) to transfer cell-type annotations from the mock reference dataset to the VA1-infected dataset **(Fig. S1C)**. At 1 dpi, VA1-infected organoids consisted of 71% progenitor cells and 29% enterocytes **(Fig. 2A-B)**. This cell type distribution differed significantly from the mock 1 dpi condition (Fisher’s exact test, p-value = 6.87 × 10^-7^) and was driven by a loss in goblet cells from 2% to 0% (proportion test, p-value = 8.50 × 10^-6^), an increase in progenitor cells from 65% to 71% (proportion test, p-value = 6.47 × 10^-4^), and a decrease in enterocytes from 33% to 29% (proportion test, p-value = 0.0140) in the VA1-infected condition. At 3 dpi, the VA1-infected organoids consisted of 56% progenitor cells, 36% enterocytes, 7% goblet cells, and 1% enteroendocrine cells **(Fig. 2C-D)**. This cell type distribution also differed significantly from the mock 3 dpi condition (Fisher’s exact test, p-value < 2.2 × 10^-16^) and was driven by a decrease in progenitor cells from 68% to 56% (proportion test, p-value < 2.2 × 10^-16^), an increase in enterocytes from 21% to 36% (proportion test, p-value < 2.2 × 10^-16)^, and a decrease in goblet cells from 10% to 7% (proportion test, p-value = 7.15 × 10^-5^) in the VA1-infected condition.

**Figure 2.**
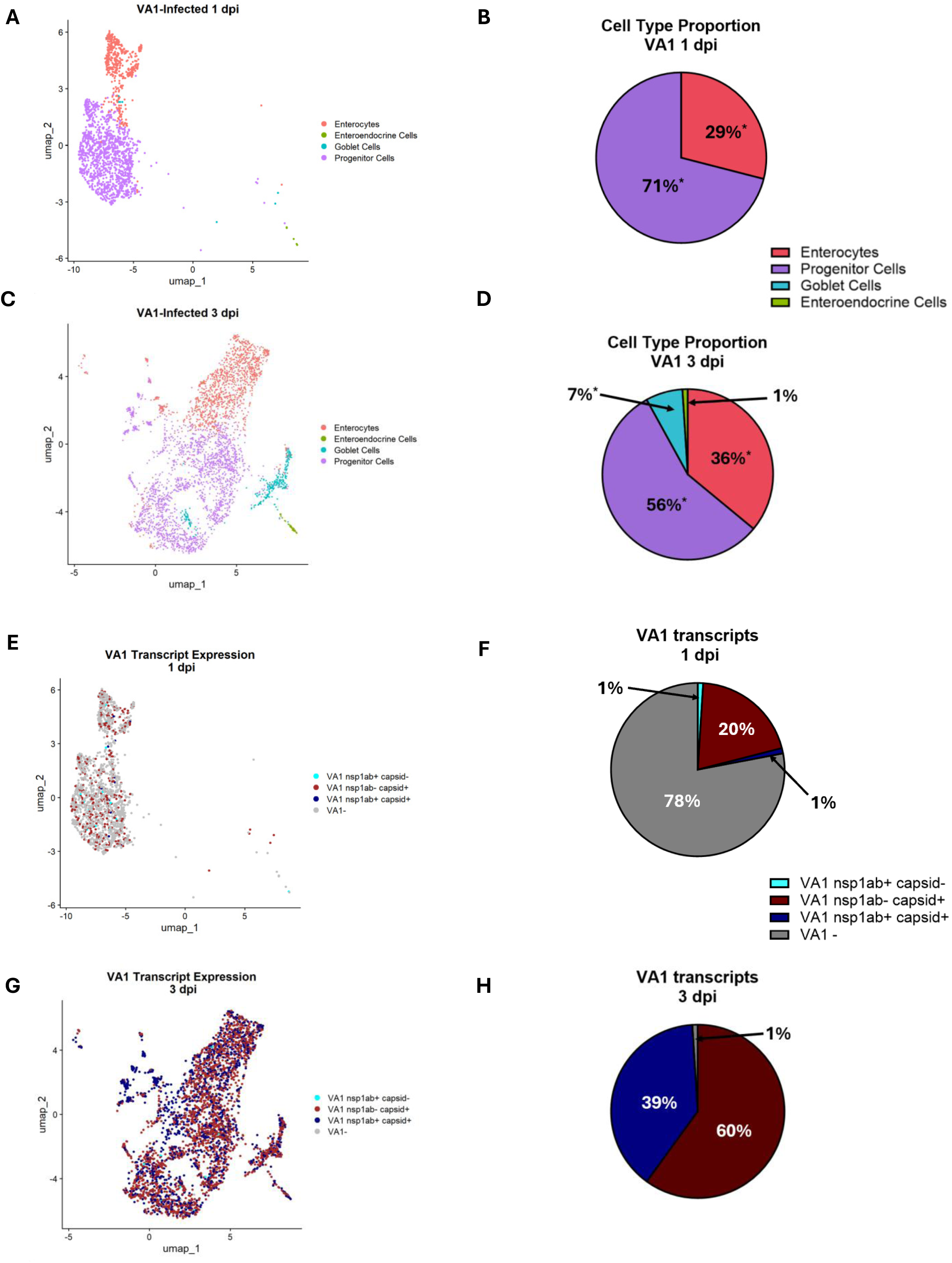
VA1 changes epithelial cell type proportions in infected human intestinal organoids. **(A)** Uniform manifold approximation and projection (UMAP) plot of VA1-infected cells at 1 dpi and **(C)** 3 dpi annotated by cell type. **(B)** Pie chart showing the distribution of cell types in the VA1-infected sample at 1 dpi and **(D)** 3 dpi. Asterisks next to the percentage of a cell type represent a significant difference, calculated by using two-sided two-sample tests for equality of proportions with continuity correction, between the proportion of that cell type in the mock- vs. VA1-infected samples. **(E)** UMAP plots of VA1-infected cells at 1 dpi and **(G)** 3 dpi annotated by their expression of VA1 transcripts: *nsp1ab* positive-*capsid* negative, *nsp1ab* negative-*capsid* positive, *nsp1ab* positive-*capsid* positive and VA1-negative (*nsp1ab* negative-*capsid* negative). **(F)** Pie chart showing the distribution of the different VA1 transcript combinations possible in the VA1-infected sample at 1 dpi and **(H)** 3 dpi.

Together, these data indicate that VA1 inoculation alters the epithelial cell type proportions in this culture. Specifically, VA1 infection decreased goblet cells at both timepoints, which contrasts with the no change in the goblet cell proportion after murine astrovirus infection despite their preferential tropism for this cell type (35). Furthermore, progenitor cells and enterocytes displayed differential effects; whereby the former decreased and the latter increased irrespective of timepoint.

To analyze VA1 transcript expression in the VA1-infected samples, we quantified cells containing VA1 nsp1ab and/or capsid transcripts. VA1 nsp1ab encodes the nonstructural polyprotein from the genomic RNA, whereas VA1 capsid encodes structural proteins expressed via the subgenomic RNA (23). At 1 dpi, VA1 transcripts were detected in both enterocytes and progenitor cells (**Fig. 2E**). At this timepoint, 22% of all cells had detectable VA1 transcripts, including 20% that were capsid-positive/nsp1ab-negative, 1% that were nsp1ab-positive/capsid-negative, and 1% that was positive for both transcripts **(Fig. 2F)**. At 3 dpi, VA1 transcripts were detected in most cells, with VA1 transcripts detected across all annotated cell types (**Fig. 2G**). At this timepoint, 99% of all cells were positive for VA1 transcripts, with 60% capsid-positive/nsp1ab-negative and 39% positive for both transcripts, while 1% contained no VA1 transcripts **(Fig. 2H)**.

Collectively, these data demonstrate that VA1 infection alters epithelial cell type proportions and that VA1 transcripts are detected in more cells of all cell types over time, increasing from 22% of cells at 1 dpi to 99% at 3 dpi.

### VA1 infection induces a cell type-specific response in enterocytes and progenitor cells

To better understand cell type-specific responses to VA1, we performed differential gene expression analysis on enterocytes and progenitor cells. Goblet and enteroendocrine cells were excluded from this analysis because their low abundance precluded statistical analysis. Differential gene expression analysis was performed between mock-infected cells and VA1 transcript-positive cells from the VA1-infected samples at both 1 and 3 dpi. Differentially expressed genes (DEGs) were defined as genes with a > 2-fold change and an adjusted p-value < 0.05. VA1 transcripts were excluded from DEG counts.

At 1 dpi, enterocytes had 354 DEGs, with 262 genes (74%) upregulated in the VA1-positive condition and 92 (26%) downregulated genes **(Fig. 3A, Fig. S2A, Table S2)**. In comparison, progenitor cells had 62 DEGs at 1 dpi, with 52 (84%) upregulated genes and 10 (16%) downregulated genes **(Fig. 3B, Fig. S2B, Table S2)**. These results indicate that enterocytes enact a broader transcriptional response to VA1 infection at 1 dpi than progenitor cells, and both cell types primarily upregulate genes in response to infection.

**Figure 3.**
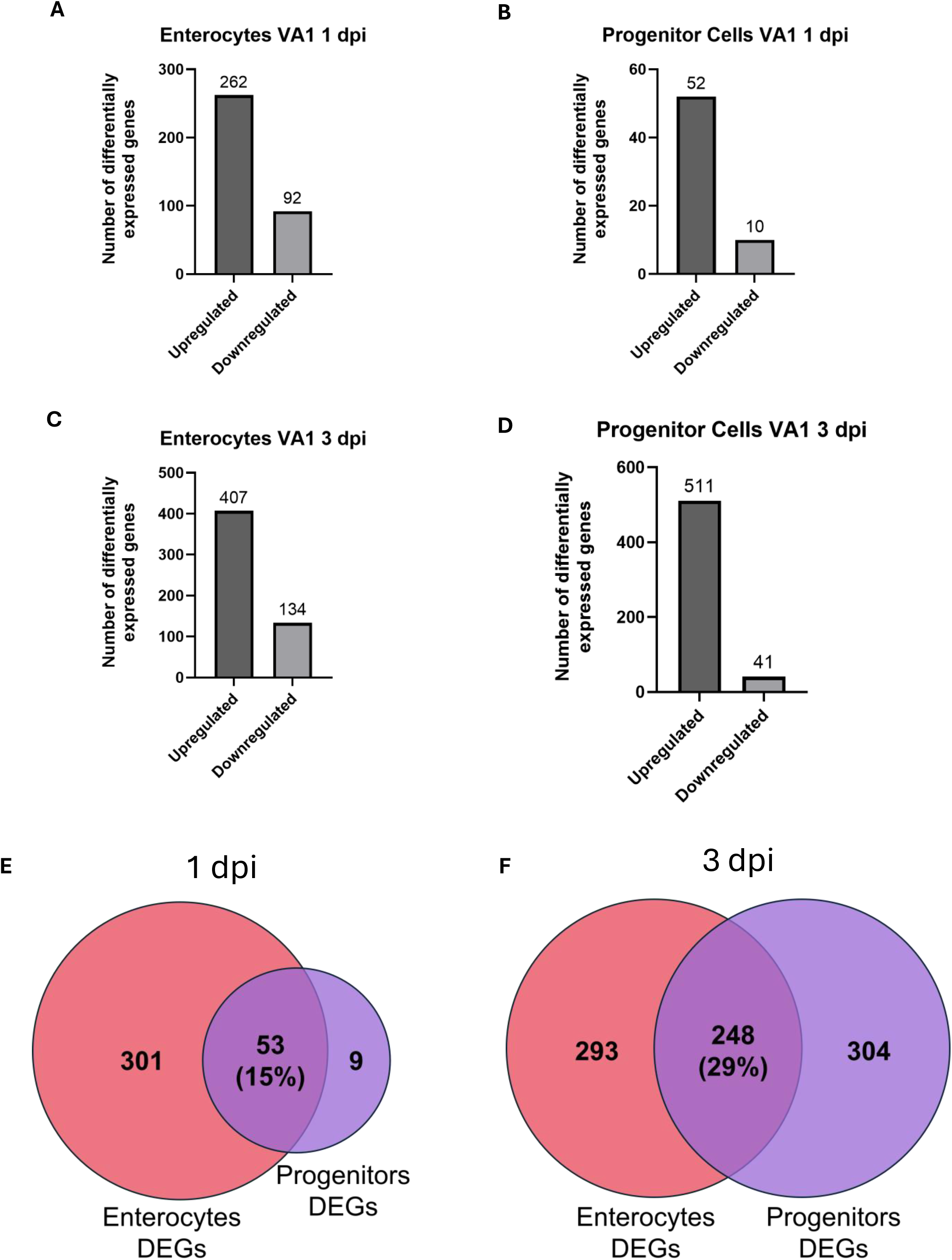
VA1 infection induces a cell type-specific response in enterocytes and progenitor cells. **(A-D)** Bar graphs showing the number of differentially expressed genes when comparing between mock-inoculated and VA1 transcript-positive enterocytes at **(A)** 1 and **(C)** 3 dpi and progenitor cells at **(B)** 1 and **(D)** 3 dpi. Differentially expressed genes were those with an adjusted p-value < 0.05 and log2 fold change larger than 1 or smaller than −1. **(E-F)** Venn diagrams showing numbers and percentages of cell type-exclusive and shared differentially expressed genes between enterocytes and progenitor cells at **(E)** 1 and **(F)** 3 dpi.

At 3 dpi, enterocytes had 541 DEGs, with 407 (75%) upregulated genes and 134 (25%) downregulated genes in the VA1-compared to mock-infected condition **(Fig. 3C, Fig. S2C, Table S2)**. Progenitor cells had 552 DEGs at 3 dpi, with 511 (93%) upregulated and 41 (7%) downregulated genes **(Fig. 3D, Fig. S2D, Table S2)**. Thus, by 3 dpi, enterocytes and progenitor cells showed a more comparable number of DEGs.

To compare the transcriptional responses of enterocytes and progenitor cells, we next identified DEGs shared between the two cell types or exclusively expressed in either cell type. At 1 dpi, 53 DEGs were shared between enterocytes and progenitor cells, representing 15% of the total 363 DEGs identified across both cell types **(Fig. 3E, Table S3)**. At this timepoint, 301 DEGs were enterocyte-exclusive and 9 DEGs were progenitor cell-exclusive **(Fig. 3E, Table S3)**. At 3 dpi, 248 DEGs were shared between enterocytes and progenitor cells, representing 29% of the total 845 DEGs identified across both cell types **(Fig. 3F, Table S3)**. At this timepoint, 293 DEGs were enterocyte-exclusive and 304 DEGs were progenitor cell-exclusive **(Fig. 3F, Table S3)**.

Taken together, these data demonstrated that enterocytes and progenitor cells share part of their transcriptional response to VA1 but also exhibit cell type-specific transcriptional changes.

### Enterocytes and progenitor cells share components of the antiviral response to VA1

To identify biological pathways associated with shared and cell type-specific responses to VA1, DEG lists were analyzed using Gene Ontology (GO) Biological Process enrichment (36–38) and transcription factor enrichment analyses (39–43). Redundant GO Biological Process terms were first reduced based on semantic similarity using the simplify function from clusterProfiler (44, 45) with a cutoff of 0.7. Among the remaining terms, a less significant term was excluded if at least 50% of its contributing DEGs overlapped with a more significant term.

At 1 dpi, shared upregulated DEGs between enterocytes and progenitor cells were enriched for biological processes associated with antiviral and stress responses. The top five non-redundant, statistically significant biological processes were “defense response to virus,” “response to hypoxia,” “response to type I interferon,” “negative regulation of viral genome replication,” and “interferon-mediated signaling pathway” **(Fig. 4A, Table S4)**. Transcription factor enrichment analysis identified interferon regulatory factor 8 (IRF8) as a significant hit among the shared upregulated DEGs **(Fig. 4B, Table S5)**. Together, these data indicate both enterocytes and progenitor cells activate a shared antiviral defense response to VA1 infection at 1 dpi.

**Figure 4.**
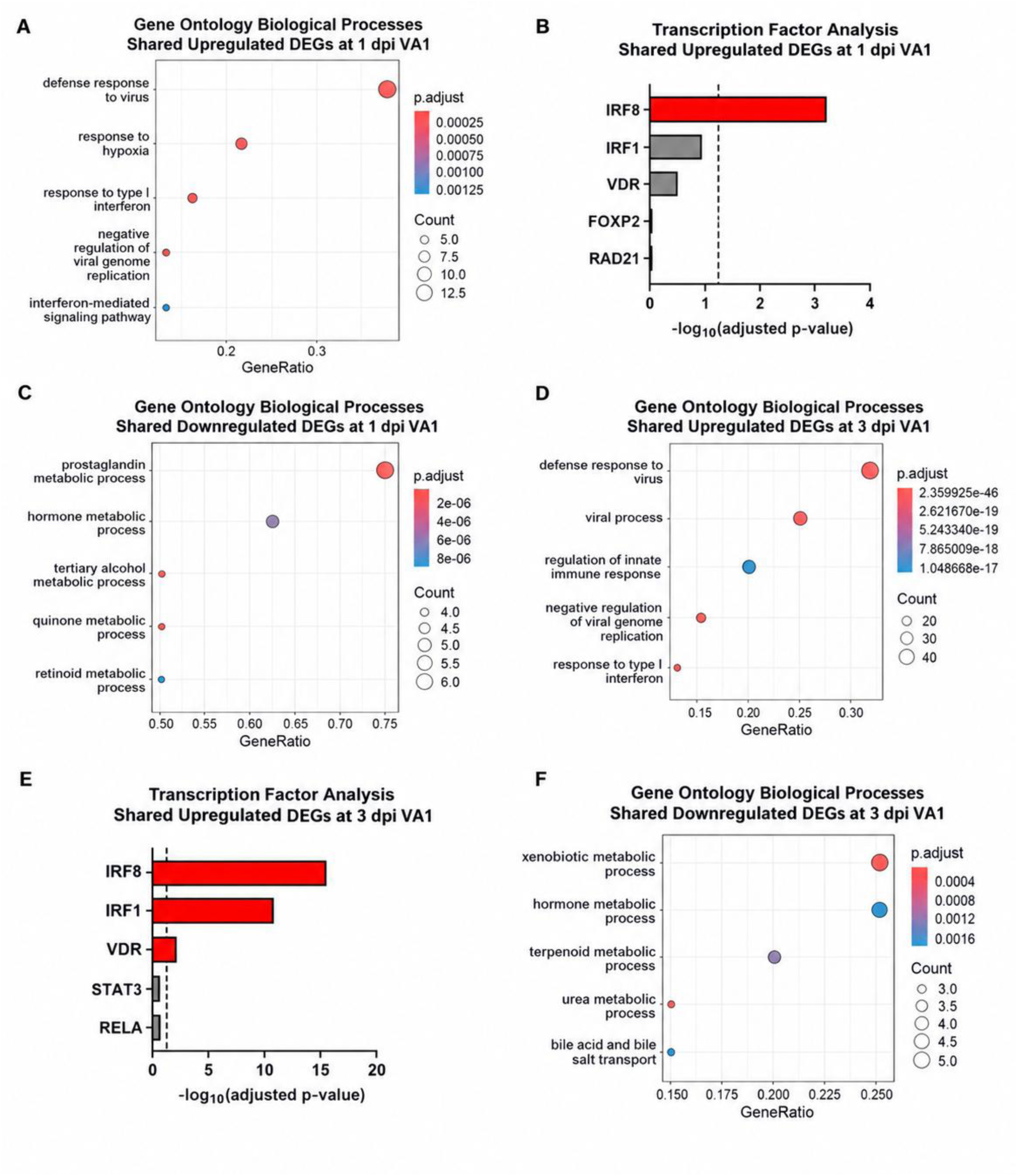
Enterocytes and progenitor cells share components of the antiviral response to VA1. **(A)** Gene Ontology Biological Processes top statistically significant hits and **(B)** transcription factor enrichment analysis top five hits for shared upregulated DEGs between enterocytes and progenitor cells at 1 dpi. **(C)** Gene Ontology Biological Processes top statistically significant hits for shared downregulated DEGs between enterocytes and progenitor cells at 1 dpi. **(D)** Gene Ontology Biological Processes top statistically significant hits and **(E)** transcription factor enrichment analysis top five hits for shared upregulated DEGs between enterocytes and progenitor cells at 3 dpi. **(F)** Gene Ontology Biological Processes top statistically significant hits for shared downregulated DEGs between enterocytes and progenitor cells at 3 dpi. Statistically significant hits were those with an adjusted p-value < 0.05. For **(B)** and **(E)**, dashed line represents statistical significance threshold: -log10 adjusted p-value > 1.30. Statistically significant transcription factor hits are shown in red.

Shared downregulated DEGs at 1 dpi were enriched for metabolic processes involving prostaglandins, hormones, tertiary alcohols, quinones, and retinoids **(Fig. 4C, Table S4)**. These enrichments were primarily driven by four aldo-keto reductase genes: AKR1B10, AKR1C1, AKR1C2 and AKR1C3. No significant transcription factor enrichment was identified among the shared downregulated DEGs at 1 dpi.

Similar to day 1, shared upregulated DEGs between enterocytes and progenitor cells at 3 dpi were enriched for biological processes related to antiviral immunity. **(Fig. 4D, Table S4)**. Transcription factor enrichment analysis again identified IRF8, as well as IRF1 and the vitamin D receptor (VDR) as significant hits **(Fig. 4E, Table S5)**, all of which function in innate immunity (46, 47).

Shared downregulated DEGs at 3 dpi were enriched for metabolic processes involving xenobiotics, hormones, terpenoids, urea, and bile acids/salts. **(Fig. 4F, Table S4)**. No significant transcription factor enrichment was identified among the 23 shared downregulated DEGs at 3 dpi.

Collectively, these data indicate that a shared antiviral response is active in both enterocytes and progenitor cells at 1 and 3 dpi, while metabolic changes suggest compromised digestive, detoxification, and regenerative capacities in both cell types at 3 dpi.

### Enterocyte-specific responses to VA1 involve early cholesterol-associated programs and broader metabolic remodeling

To identify enterocyte-specific responses to VA1 infection, enterocyte-exclusive DEGs were analyzed using Gene Ontology Biological Process and transcription factor enrichment analyses.

At 1 dpi, enterocyte-exclusive upregulated DEGs were enriched for two major categories of biological processes: antiviral responses and cholesterol-associated metabolism. Antiviral terms included “response to virus,” “viral process,” and “negative regulation of viral genome replication,” whereas cholesterol-associated terms included “cholesterol metabolic process” and “sterol biosynthetic process” **(Fig. 5A, Table S6)**. Although antiviral terms were also enriched among DEGs shared by enterocytes and progenitor cells, the enterocyte-exclusive DEGs expanded this shared response by including additional genes involved in viral sensing and signaling, such as Z-DNA-binding protein 1 (ZBP1) and DExD/H-box helicase 60 (DDX60), as well as antiviral effector genes, including radical S-adenosyl methionine domain-containing 2/viperin (RSAD2) and 2′-5′-oligoadenylate synthetase 2 (OAS2) **(Table S3)**. Thus, enterocytes activated an expanded antiviral gene repertoire alongside a cholesterol-associated transcriptional program at 1 dpi. Transcription factor enrichment analysis of the enterocyte-exclusive upregulated DEGs identified IRF1 and IRF8, VDR, and GATA-binding proteins 1 and 2 (GATA1 and GATA2) as significant hits **(Fig. 5B, Table S7)**.

**Figure 5.**
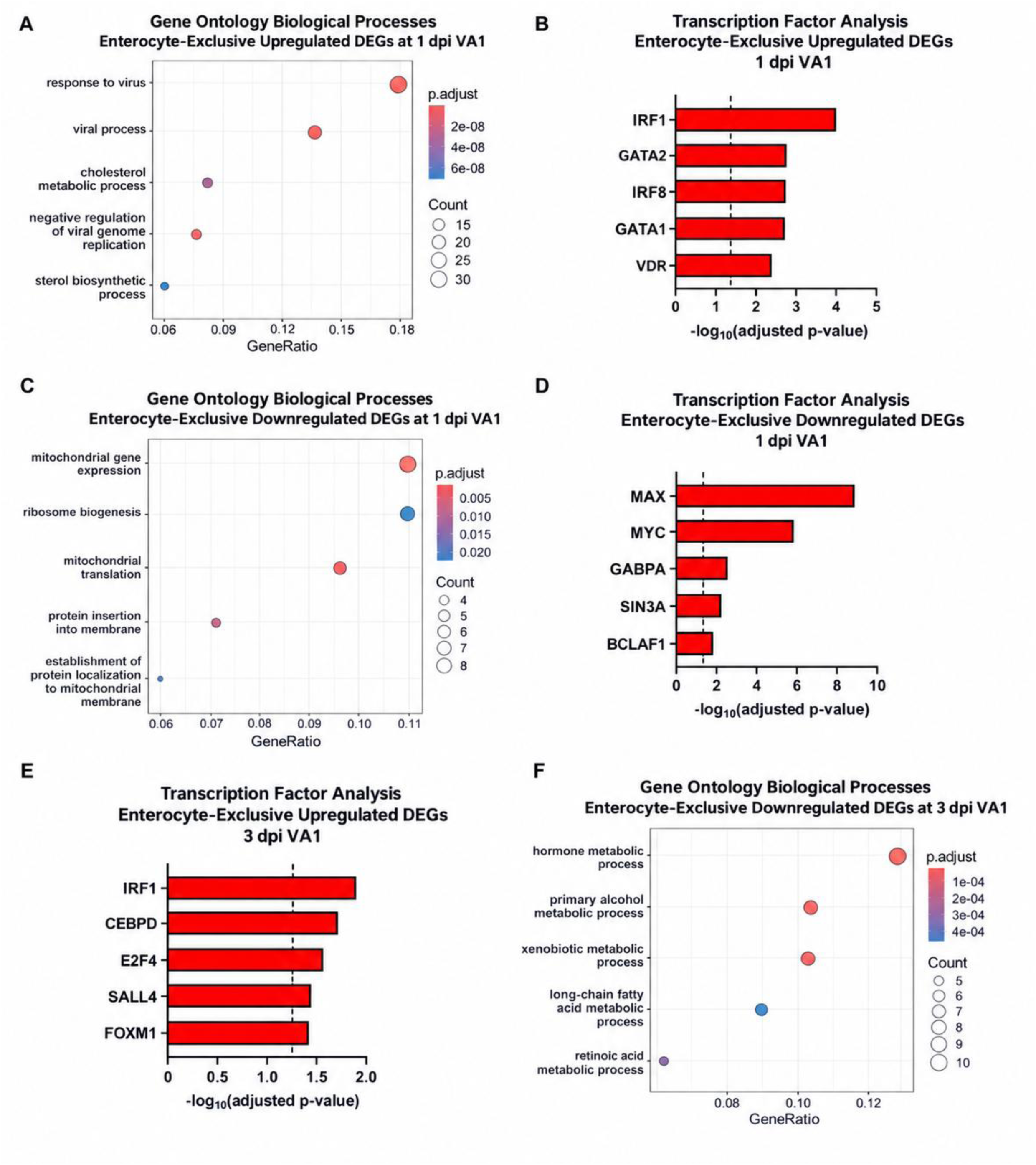
Enterocyte-specific responses to VA1 involve early cholesterol-associated programs and broader metabolic remodeling. **(A)** Gene Ontology Biological Processes top statistically significant hits and **(B)** transcription factor enrichment analysis top five hits for enterocyte-exclusive upregulated DEGs at 1 dpi. **(C)** Gene Ontology Biological Processes top statistically significant hits and **(D)** transcription factor enrichment analysis top five hits for enterocyte-exclusive downregulated DEGs at 1 dpi. **(E)** Transcription factor enrichment analysis top five hits for enterocyte-exclusive upregulated DEGs at 3 dpi. **(F)** Gene Ontology Biological Processes top statistically significant hits for enterocyte-exclusive downregulated DEGs at 3 dpi. Statistically significant hits were those with an adjusted p-value < 0.05. For **(B)**, **(D)** and **(E)** dashed line represents statistical significance threshold: -log10 adjusted p-value > 1.30. Statistically significant transcription factor hits are shown in red.

Enterocyte-exclusive downregulated DEGs at 1 dpi were enriched for biological processes related to mitochondrial function and protein synthesis, including “mitochondrial gene expression,” “ribosome biogenesis,” “mitochondrial translation,” “protein insertion into membrane,” and “establishment of protein localization to mitochondrial membrane” **(Fig. 5C, Table S6).** Transcription factor enrichment analysis of these downregulated DEGs identified MYC, MYC-associated factor X (MAX), GA-binding protein alpha (GABPA), Switch-Independent 3A (SIN3A), and Bcl-2–associated transcription factor 1 (BCLAF1) as significant hits **(Fig. 5D, Table S7)**. These results indicated that VA1 infection of enterocytes may be associated with downregulation of genes involved in mitochondrial processes and protein translation, along with genes downstream of cell growth and stress response regulators at 1 dpi (48, 49).

At 3 dpi, enterocyte-exclusive upregulated DEGs did not yield significantly enriched Gene Ontology Biological Process terms. However, transcription factor enrichment analysis identified IRF1, CCAAT/enhancer-binding protein delta (CEBPD), E2F transcription factor 4 (E2F4), spalt-like transcription factor 4 (SALL4), and forkhead box M1 (FOXM1) as significant hits **(Fig. 5E, Table S7)**. These transcription factors have established roles in innate immune and inflammatory signaling (IRF1 and CEBPD), cell-cycle regulation (E2F4 and FOXM1), and the maintenance of stemness or plasticity of cells (SALL4) (46, 50–53).

Enterocyte-exclusive downregulated DEGs at 3 dpi were enriched for metabolic processes involving hormones, primary alcohols, xenobiotics, long chain fatty acids, and retinoic acids **(Fig. 5F, Table S6)**. These enrichments overlapped partially with metabolic pathways identified among enterocyte/progenitor shared downregulated DEGs at the same timepoint (see **Fig. 4F**), indicating that VA1-associated metabolic changes are detected broadly but are especially evident in enterocytes. No significant transcription factor enrichment was identified among the enterocyte-exclusive downregulated DEGs at 3 dpi.

Taken together, these data identify an enterocyte-specific response to VA1 at 1 dpi characterized by activation of a cholesterol-associated transcriptional program and an expanded antiviral gene repertoire beyond the core response shared with progenitor cells, while coinciding with downregulation of mitochondrial and biosynthetic processes. At 3dpi, enterocytes exhibited broader metabolic downregulation, along with transcription factor enrichment associated with immune signaling and cell-state regulation. Thus, the enterocyte transcriptional response transitioned from an early antiviral response and remodeling of cholesterol homeostasis to extensive metabolic modulation at 3 dpi.

### Progenitor-exclusive DEGs reveal immune and epithelial-defense signatures at 3 dpi

To characterize progenitor cell-specific responses to VA1, progenitor-exclusive DEGs were analyzed using Gene Ontology Biological Process and transcription factor enrichment analyses.

At 1 dpi, only nine progenitor-exclusive DEGs were identified, including seven upregulated and two downregulated genes. The upregulated DEGs did not yield any significantly enriched Gene Ontology Biological Process terms. Analysis of the two downregulated DEGs identified enrichment for “disaccharide metabolic process” and “cellular glucuronidation” **(Fig. S3A, Table S6)**. However, each enrichment was based on a single gene expressed in only a small proportion of VA1-positive progenitor cells. No significant transcription factor enrichment was identified among the progenitor-exclusive upregulated or downregulated DEGs at 1 dpi.

At 3 dpi, progenitor-exclusive upregulated DEGs did not yield any significantly enriched Gene Ontology Biological Process terms. Nevertheless, inspection of the most highly upregulated genes suggested inflammatory and epithelial-defense responses. These genes included the chemokines *CCL5*, *CXCL10*, and *CXCL11*; reactive oxygen species-associated antimicrobial defense genes *DUOX2* and *DUOXA2*; and additional immune or mucosal defense genes, including *IL32*, *ITLN1*, *SOCS1*, *TNFAIP2*, and *IFITM10*. Several of these genes were detected in only a small proportion of VA1-positive progenitor cells, suggesting that this signature may be concentrated within a subset of the progenitor-cell population. Progenitor-exclusive downregulated DEGs were enriched for “positive regulation of ATP metabolic process” **(Fig. S3B, Table S6)** but otherwise lacked a clear shared functional pattern. No significant transcription factor enrichment was identified among the progenitor-exclusive upregulated or downregulated DEGs at 3 dpi.

Overall, progenitor cells showed limited evidence of a functional response distinct from enterocytes at 1 dpi. At 3 dpi, however, progenitor-exclusive DEGs suggested inflammatory and epithelial-defense responses, although these genes did not converge on significantly enriched biological processes or transcription factor-associated programs.

### Cholesterol synthesis is required for optimal VA1 replication in T84 and Caco-2 cells

To further investigate the role of cholesterol homeostasis in VA1 replication in enterocytes, we first tested whether availability of extracellular cholesterol affected VA1 replication in T84 and Caco-2 cells, human colorectal carcinoma cell lines commonly used in the field. These cells were cultured in regular or lipid-depleted medium and inoculated with VA1 at a multiplicity of infection (MOI) of 100 genome copies per cell (∼0.01 focus-forming units [FFU] per cell). VA1 genome copies increased from 0 to 1 dpi in T84 cells and from 0 to 2 dpi in Caco-2 cells under both lipid-replete and lipid-depleted conditions, confirming successful infection **(Fig. 6A–B)**. However, VA1 genome copies did not differ between conditions at either 0 or 1 dpi in T84 cells **(Fig. 6A)**. In contrast, extracellular lipid depletion significantly increased titers at 0 dpi in Caco-2 cells, although titers were similar between conditions at 2 dpi **(Fig, 6B)**. These findings suggest that extracellular lipid depletion enhances VA1 binding to Caco-2 but not T84 cells. However, the convergence of Caco-2 titers by 2 dpi despite a higher titer at 0 dpi under lipid-depleted conditions suggests that lipid depletion may also have a minor impact on VA1 replication.

**Figure 6.**
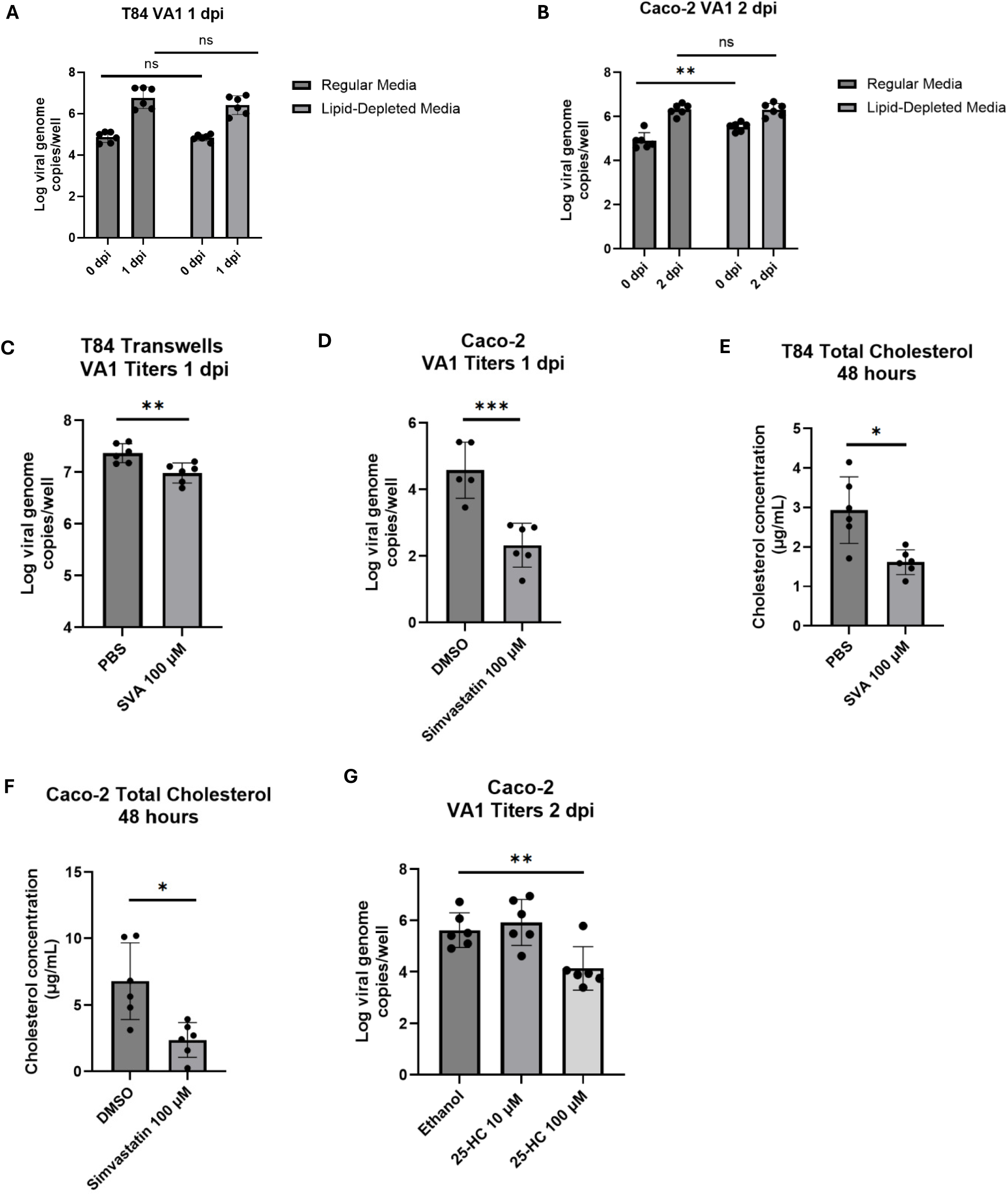
Cholesterol biosynthesis inhibition and 25-hydroxycholesterol affects VA1 replication in T84 and Caco-2 cells. **(A)** T84 and **(B)** Caco-2 cells were inoculated with VA1 at an MOI of 100 for an hour after a 24h pre-treatment with lipid-depleted or regular media. Cells were collected at 0 and 1 dpi (for T84) or 2 dpi (for Caco-2) and VA1 genome copies were measured through qRT-PCR. Statistical analysis was performed using 2-way ANOVA tests. **(C)** T84 transwells and **(D)** Caco-2 cells were inoculated with VA1 at an MOI of 1000 (for T84) or 100 (for Caco-2) for an hour after a 24h pre-treatment with indicated concentrations of simvastatin with their respective vehicle controls. Cells were collected at 1 dpi and VA1 genome copies were measured through qRT-PCR. Statistical analysis was performed using two-tailed students-t tests. **(E)** T84 Transwells and **(F)** Caco-2 cells were treated with indicated concentrations of SVA (for T84), simvastatin (for Caco-2), or the respective vehicle controls for 48 hours. Total cholesterol levels were measured using the Amplex Red Cholesterol Assay Kit. Statistical analysis was perfomed using Welch’s t-tests. **(G)** Caco-2 cells were inoculated with VA1 at an MOI of 100 for an hour. Media after infection contained indicated concentrations of 25-HC. Cells were collected at 2 dpi and VA1 genome copies were measured through qRT-PCR. Statistical analysis was performed using two-tailed students-t tests. Experiments represent combined data from at least two independent experiments with three technical replicates each. ***, *p <*0.001; **, *p* <0.01; *, *p <*0.05; ns, not significant.

We next examined whether pharmacologic inhibition of cholesterol biosynthesis affects VA1 replication. To better mimic the polarized intestinal epithelium, T84 cells were seeded on transwell inserts for four days until they reached an average transepithelial electrical resistance of ∼2,000 Ω*cm^2^, which is above the 1,000 Ω*cm^2^ threshold for establishment of a tight intestinal epithelial barrier in T84 transwells (54). Cells were then pre-treated for 24 hours with 100 µM simvastatin sodium salt (SVA), the active, acid form of the inhibitor of 3-hydroxy-3-methylglutaryl-CoA reductase (HMGCR), the rate-limiting enzyme of the cholesterol biosynthesis pathway (55). Next, cells were inoculated with VA1 at an MOI of 1000 genome copies/cell (∼0.1 FFU/cell). SVA treatment resulted in a statistically significant decrease in VA1 genome copies at 1 dpi compared to the vehicle (phosphate-buffered saline, PBS)-treated control **(Fig. 6C)**. A similar finding was made in Caco-2 cells pre-treated with 100 µM of the lactone prodrug form of simvastatin and inoculated with VA1 at an MOI of 100 genome copies (∼0.01 FFU)/cell **(Fig. 6D)** (55). Fluorometric assays confirmed that simvastatin treatment reduced total cellular cholesterol levels in both cell lines **(Fig. 6E-F)**, without reducing cell viability below 80% compared to untreated cells under the conditions tested **(Fig. S4A-B)**. These data indicate that inhibiting cholesterol synthesis reduces VA1 replication.

Another important arm of cellular cholesterol homeostasis is intracellular transport between cellular compartments. To determine whether disrupting intracellular cholesterol trafficking affected VA1 replication, T84 cells were treated with U18666A, which inhibits the NPC intracellular cholesterol transporter 1 (NPC1)-mediated transport of cholesterol out of late endosomes and lysosomes (56). Toxicity studies indicated that concentrations of up to 25 µM U18666A maintained cell viability **(Fig. S4C)**. Treatment with 10, 17.5, or 25 µM U18666A did not significantly alter VA1 genome copy numbers at 1 dpi **(Fig. S4D)**, despite immunofluorescence imaging demonstrating increased intracellular filipin-positive puncta, indicative of cholesterol accumulation in cells treated with 25 µM U18666A compared to vehicle-treated controls **(Fig. S4E–G)**. These data suggested that intracellular cholesterol transport does not play a major role during VA1 replication.

An important regulator of cholesterol homeostasis is the oxysterol 25-hydroxycholesterol (25-HC). 25-HC is produced by the enzyme cholesterol 25-hydroxylase (CH25H) and is an important negative regulator of cholesterol homeostasis. 25-HC has antiviral properties against multiple viruses (57, 58). Furthermore, CH25H in addition to its enzymatic function is also an ISG that was significantly upregulated in VA1-infected T84 transwells at 3 dpi (54). Thus, we also tested the effect of 25-HC on VA1 replication. Caco-2 cells were treated with 10 and 100 μM 25-HC for 24 hours after inoculation with VA1 at an MOI of 100 genome copies/cell (∼0.01 FFU/cell). Treatment with 25-HC significantly reduced VA1 genome copies at 2 dpi compared with the ethanol vehicle control **(Fig. 6G)**, while cell viability was not significantly reduced at these concentrations **(Fig. S4H)**. These findings demonstrate that 25-HC has antiviral activity against VA1 similar to other viruses, including the enteric noroviruses (59, 60).

Taken together, our experimental validation indicated that inhibiting cholesterol biosynthesis and treatment with 25-HC reduced VA1 replication in intestinal epithelial cells, whereas disrupting intracellular cholesterol transport had no effect. In addition, reducing extracellular lipid availability may have cell-line-specific effects on VA1 binding and replication.

## Discussion

HAstVs are understudied viruses, without approved antivirals or vaccines. They are an important cause of diarrhea in children and extraintestinal disease in immunocompromised patients, with many of the encephalitis cases being associated with the non-classical strain VA1. In this study, single-cell RNA sequencing of VA1-infected fetal ileum-derived organoids revealed shared and distinct transcriptional responses between enterocytes and progenitor cells, consistent with previously published studies showing that VA1 induced a strong IFN response and that different cell types can have different transcriptional profiles after viral infection (28, 29). One notable finding of our work was the enterocyte-specific upregulation of genes involved in cholesterol homeostasis at 1 dpi.

Subsequent experimental validation showed that simvastatin and 25-hydroxycholesterol reduced VA1 replication. On the other hand, extracellular lipid depletion had cell-line-specific effects, while blocking cholesterol transport with U18666A had no effect on VA1 genome titers. Collectively our study showed that intestinal epithelial cell types active different transcriptional programs in response to VA1 infection and identified cholesterol-related pathways as a potential target for future antiviral development.

One striking finding in our study was the detection of VA1 transcripts in all epithelial cell types at 3 dpi, highlighting that transcript detection alone does not distinguish between productive infection, abortive infection, or viral RNA uptake. This raises important questions about the relationship between viral RNA detection and productive infection. As anticipated, VA1 transcripts were detected in a subset of cells, including enterocytes and progenitor cells at 1 dpi. However, by 3 dpi, nearly all cells contained detectable VA1 transcripts. Despite this widespread detection, capsid, but not nsp1ab transcript expression, showed a bimodal distribution at 3 dpi, with most cells exhibiting relatively low expression and a smaller population, comprising approximately 12.5% of cells, exhibiting high expression **(Fig. S5A-B)**. This distribution suggests that VA1 capsid transcript-positive cells may represent biologically distinct populations with different levels of viral RNA.

Interestingly, the proportion of cells with high VA1 capsid transcript expression was within the approximately 5–20% range of cells previously reported to be positive for double-stranded RNA, a marker of active viral replication, in VA1-infected organoids (28). This raises the possibility that this VA1 capsid high expressing population represents cells supporting the complete viral life cycle and productive infection, since large numbers of capsid proteins are needed to encapsidate viral genomes and assemble. Cells with low transcript levels on the other hand may contain viral RNA without forming infectious particles, therefore representing abortive infection. One potential explanation for the wide-spread presence of viral RNA in most cells at the later timepoint is their transfer between cells through extracellular vesicles or tunneling nanotubes, as has been reported for viral RNAs in epithelial cells infected with enterovirus, influenza A virus, and porcine reproductive and respiratory syndrome virus (61, 62). Therefore, VA1 transcript positivity is useful for identifying cells exposed to or containing viral RNA but should not be interpreted as evidence of productive infection. Instead, the VA1 transcriptional signatures likely represent a mixture of cells representing productive and abortive infection. Similar observations and conclusions were made in a recent single-cell RNAseq study of human norovirus-infected duodenal organoids (63).

Another important finding was that VA1 infection altered cell type proportions in infected organoids. The differences in cell type proportions observed between VA1- and mock- inoculated samples varied over time, with a lower enterocyte proportion at 1 dpi but a higher proportion at 3 dpi, accompanied by reciprocal changes in progenitor-cell representation. Altered cell proportions were also observed following organoid infections with other enteric viruses. Specifically, single-cell analysis of HAstV1-infected organoids showed increased expression of proliferative cells as measured by the proliferation marker MKI67 and decreased enterocyte abundance (29), while rotavirus infection of intestinal organoids increased enterocyte abundance and decreased stem cell representation (64). However, no changes in cell type proportions were observed in SARS-CoV-2-infected intestinal organoids (65). These comparisons suggest that infection-associated epithelial remodeling may differ among viruses and across different timepoints of infection. Future studies are needed to investigate the mechanism(s) by which VA1 induces these changes in cell type proportions.

A further notable finding was that both enterocytes and progenitor cells mounted shared responses to VA1, including antiviral, hypoxia and metabolism related gene signatures. The upregulation of antiviral defense responses, including IFN signaling and innate immune regulation, as well as activation of transcription factors IRF8 and IRF1, at both timepoints are consistent with previous reports that the intestinal epithelium can mount strong antiviral responses against enteric viruses (28, 30, 63). In addition, hypoxia-associated pathways were also upregulated in both cell types at 1 dpi. This is particularly interesting given the multifaceted role of hypoxia in regulating innate immunity in the context of viral infections. While hypoxia has an immunosuppressive effect on Zika virus infection in monocytes by decreasing the type I IFN response, it has a pro-inflammatory effect on vesicular stomatitis virus infection in kidney cells by increasing IFN induction (66). Notably, the upregulation of hypoxia-related genes during VA1 infection contrasts with findings from human norovirus infection of the same human intestinal ileal organoid line, where hypoxia-related genes were downregulated (32). Additionally, hypoxic conditions have been shown to be proviral in the context of rotavirus infection, where hypoxia led to a decreased induction of the type III IFN response (67). Since this response is also upregulated during VA1 infection, this may suggest a proviral role for the upregulation of hypoxia-related genes following VA1 infection. Additional experiments to determine the role of hypoxia in VA1 infection are warranted in the future.

In addition to these shared upregulated pathways, VA1 infection was further associated with a downregulation in metabolic gene expression in enterocytes and progenitor cells. At 1 dpi, shared downregulated genes were enriched for metabolicprocesses involving prostaglandins, hormones, quinones, retinoids, and tertiary alcohols, driven largely by aldo-keto reductase genes, while at 3 dpi, the downregulated metabolic pathways included xenobiotics, hormones, terpenoids, urea, and bile acid/bile salt transport processes. These results suggest that VA1 infection may suppress or remodel epithelial metabolic functions over time. The downregulation of prostaglandin-related pathways is interesting because prostaglandin signaling has antiviral effects in intestinal epithelial cells in the context of HIV-1 infection (68, 69), which may suggest that a VA1-induced downregulation of these genes might help the virus to escape their antiviral function. Similarly, retinoic acids and bile acids have been shown to have antiviral roles in the context of murine norovirus, rotavirus, and porcine epidemic diarrhea virus infections (70–73), thus their downregulation by VA1 may be a strategy to evade the antiviral effects.

Taken together, these findings indicate that enterocytes and progenitor cells mount a sustained shared antiviral response to VA1, while concurrent hypoxia-associated and metabolic changes may modulate epithelial antiviral defenses and the cellular environment in which VA1 replicates. However, future studies are needed to determine whether these changes restrict or promote VA1 replication.

Beyond these shared responses, VA1 infection induced cell type specific responses, a finding also reported for HAstV-1-infected organoids and in murine astrovirus (MuAstV)-infected small intestinal tissue (29, 74). Intestinal epithelial populations have distinct baseline transcriptional profiles and functions that may influence their responses to infection (75). In the case of classical HAstV-1, single-cell analysis of infected intestinal organoids identified distinct baseline and infection-associated ISG expression profiles across cell types, including extensive transcriptional changes in enterocytes. In the case of VA1, enterocytes exhibited a broader early antiviral gene repertoire than progenitor cells during infection. These similarities suggest that enterocytes contribute prominently to infection-associated antiviral responses. This is consistent with the induction of IFN-λ in enterocytes by MuAstV, which plays an important role in protection against MuAstV infection and other enteric viruses (74, 76).

Additionally, both VA1- and norovirus-infected enterocytes in intestinal organoids upregulated IL15 at 3 and 2 dpi, respectively. This may suggest a shared epithelial response with the potential to modulate local immune activity and promote repair since IL15 on enterocytes promotes survival, maintenance and maturation of intraepithelial lymphocytes and activates intracellular signaling cascades to promote proliferation and survival of intestinal epithelial cells, aiding in mucosal turnover and barrier maintenance (77–79).

Concurrent upregulation of IL15RA, the receptor used by intestinal epithelial cells to *trans*-present IL15 to neighboring cells (80), in VA1-infected enterocytes at 3 dpi further suggests increased capacity for IL-15 *trans*-presentation, which could support the survival and activation of intraepithelial lymphocytes. However, functional studies are needed in the future to determine whether these transcriptional changes translate into enhanced IL-15 signaling in intestinal organoids.

Conversely, the limited number of progenitor-exclusive antiviral DEGs at 1 dpi may partly reflect the higher basal ISG expression previously reported in intestinal stem cells compared with enterocytes and goblet cells (29). Stem cells from several other tissues also constitutively express subsets of antiviral genes, providing an intrinsic antiviral state before infection (81). Together, these findings further highlight that different cell types mount cell-type-specific responses to viral infections.

Another key finding was the enterocyte-exclusive upregulation of genes involved in cholesterol homeostasis at 1 dpi. Cholesterol plays an important role in the life cycles of many positive-sense single-stranded RNA viruses. For example, cholesterol-rich lipid rafts contribute to the entry of other enteric viruses, including norovirus and rotavirus (82–84). Cholesterol also contributes to the formation of replication organelles in positive-sense RNA viruses. For example, hepatitis C virus replicates in cholesterol-enriched double-membrane vesicles (85). HAstVs have similarly been shown to depend on virus-induced membrane rearrangements for efficient replication, suggesting a role for cholesterol in their replication (86). Our finding that treatment with simvastatin, which inhibits the rate-limiting enzyme in the cholesterol biosynthesis pathway, significantly reduced VA1 genome titers, supports the critical role for cholesterol during HAstV infection. However, the stage of the VA1 life cycle blocked by simvastatin treatment remains to be investigated and future studies defining the role of cholesterol in the VA1 life cycle are needed. Additionally, our finding that treatment with 25-HC, a cholesterol-regulating oxysterol, reduced VA1 replication is consistent with previously published studies where 25-HC had antiviral effects against a variety of viruses (57, 58). However, the mechanism behind its antiviral effect against VA1 requires further studies. More broadly, whether dependence on cholesterol synthesis is a conserved feature of HAstV biology remains to be determined.

This study also has several limitations. First, single-cell RNA sequencing detects viral transcripts but cannot independently distinguish productive infection from abortive infection, viral RNA uptake, or paracrine responses to infected neighboring cells. Our previous work also identified progenitor cells, enterocytes and goblet cells as VA1-infected cell types, specifically showing VA1 dsRNA replication intermediates and capsid protein production by flow cytometry and immunofluorescence, respectively (28). Therefore, the combination of approaches demonstrates the multi-cellular tropism of VA1 for progenitor cells, enterocytes and goblet cells in HIOs. Future work will be needed to verify this cellular tropism in infected individuals and to clarify a potential tropism for enteroendocrine cells that was indicated by the presence of viral RNA in our single cell RNAseq study. Second, the low abundance of goblet and enteroendocrine cells limited our ability to analyze their transcriptional responses, even though these lineages may play important roles in intestinal defense and epithelial function. To investigate goblet- and enteroendocrine cell-specific responses, future studies could culture HIOs under lineage-specific differentiation conditions before viral infection, using Notch inhibition with the γ-secretase inhibitor dibenzazepine (DBZ) to enrich goblet cells or combined Wnt, Notch, and MAPK pathway inhibition to enrich enteroendocrine cells (87, 88). Third, fetal ileum-derived organoids provide a valuable model of human intestinal epithelium but do not fully capture the complexity of the intestinal environment, including immune cells, microbiota, stromal cells, and luminal factors such as bile acids and dietary metabolites. The microbial metabolite butyrate was recently shown to reverse VA1-induced barrier permeability in T84 cells (54). Therefore, future studies of HAstV infections in more complex HIO co-cultures are needed to elucidate how these factors impact the cell type specific and shared responses of the intestinal epithelium. Fourth, because the dataset includes limited timepoints, it may miss earlier entry-associated events or later outcomes related to epithelial repair, persistence, or cell death. Analyzing single-cell transcriptomics of VA1-infected HIOs at early timepoints, such as 6 or 12 hours post-infection, or later timepoints, such as 4 and 5 dpi, once viral titers have plateaued (28), would provide a broader understanding of the kinetics of VA1 infection and its time-dependent effects in the intestinal epithelium. Finally, the transcriptomic dataset was derived from a single donor, limiting our ability to account for the effects of genetic variation and donor age on responses to infection. Donor age may be particularly important because intestinal organoids derived from younger individuals have been shown to mount weaker innate immune responses than those derived from adults (89). Therefore, repeating experiments in additional organoid lines derived from different donors, including donors of different ages, could provide information on the role of host genetics and donor age on VA1 pathogenesis.

Overall, this study demonstrates that VA1 infection induces both shared and cell type-specific transcriptional responses in human fetal ileum-derived organoids. While enterocytes and progenitor cells both activate antiviral IFN-associated pathways, enterocytes exhibit a broader and more metabolically distinct response, including regulation of cholesterol homeostasis genes at early times during infection. These findings suggest that differentiated epithelial cells may play a particularly important role in shaping the intestinal response to VA1 and identify cholesterol metabolism, especially cholesterol synthesis, as a promising pathway for further mechanistic investigations. By defining how individual epithelial cell types respond to VA1 infection, this work provides new insight into HAstV biology and highlights host pathways that may be exploited for future antiviral strategies.

## Methods

### Viruses and cells

HAstV-VA1 (passage #3) was expanded in Caco-2 cells to create a virus stock as described before (28). Mock Caco-2 lysates were produced through the same process. Human intestinal organoid (HIO) line HT-124-I was derived from fetal ileum and obtained from the University of Michigan Medical School Translational Tissue Modeling Laboratory. Organoids were maintained in a complete media with growth factors (CMGF+) as described in (32). HIOs were cultured in 6-well plates (CytoOne CC7682) embedded in Matrigel (Corning 356231) with a total of 20 domes per well. Media was changed every day, and the HIOs were passaged every 7 days.

T84 cells were cultured in Advanced DMEM/F12 supplemented with 10% heat-inactivated fetal bovine serum (FBS) (HyClone #SH30396.03), 1% penicillin–streptomycin, 1% HEPES, and 1% L-glutamine (Gibco 25030–081). Caco-2 cells were cultured in DMEM-10 prepared using Gibco DMEM medium (Cat. No. 11995-065) with 4.5 g/L D-glucose and 110 mg/L sodium pyruvate, supplemented with 10% heat-inactivated fetal bovine serum, 1% penicillin–streptomycin, 1% L-glutamine, 1% HEPES, and 1% MEM non-essential amino acids (Gibco 11140–050). Cells were maintained in tissue-culture-treated flasks at 37°C in a humidified atmosphere containing 5% CO₂.

### VA1 infections of human intestinal organoids for single-cell RNA sequencing (scRNAseq) sample submission

FI124 HIOs were embedded in Matrigel and grown in CMGF+ media for five days until HIO spheroids had reached maximum growth as before (28). Healthy HIO spheroids were then dissociated into cell aggregates by pipetting and passing through a 25G x 5/8” needle. The cell aggregates obtained were washed, resuspended in 0.5 ml CMGF+ and split into 2 groups of equal volume. The mock-infected groups were mixed with 0.25 ml Caco-2 cell lysate while the VA1-infected groups were mixed with the same volume of virus stock containing ≈ 1 × 10^7^ genome copies of VA1. HIO infection was allowed to proceed for 1 hour at 37°C, after which cells were washed. Mock-treated HIO and VA1-infected HIO cell aggregates were embedded in Matrigel and grown in CMGF+ into healthy spheroids as before. After confirming infection by qRT-PCR from cell fractions at 1 and 3dpi, cells were collected for scRNAseq sample preparation.

### Single-cell dissociation and library preparation for scRNA-seq

Mock-infected and VA1-infected HIO spheroids were collected at 1 and 3 dpi to prepare single-cell suspensions. One biological replicate was collected per condition and time point. Each replicate consisted of 16 Matrigel domes, which were pooled before library preparation. The 1 and 3 dpi samples were obtained from independent infection experiments. A total of four sequencing libraries—Mock 1 dpi, VA1 1 dpi, Mock 3 dpi, and VA1 3 dpi were prepared separately.

HIO spheroids were suspended in PBS containing 0.5 mM ethylenediaminetetraacetic acid (EDTA) and dissociated into cell aggregates by pipetting and passage through a 25-gauge × 5/8-inch needle. HIO cell aggregates were collected by centrifugation, resuspended in TrypLE Express Enzyme (Gibco 12604013), incubated at 37 °C for 3 minutes, and pipetted up and down to generate single-cell suspensions. From this point onward, cells were maintained on ice. Cells were passed through a 40 µm cell strainer to remove remaining aggregates and collected by centrifugation at 500 × g for 5 minutes at 4 °C.

Cells were resuspended in Advanced DMEM/F-12 supplemented with 10% fetal bovine a target concentration of 1 × 10^6^ cells/mL. Cell viability, measured by the LUNA Fx7 Automated Cell Counter (Logos Biosystems) at 1 dpi and the Countess II Automated Cell Counter (Invitrogen) at 3 dpi, was greater than 75% for all samples. Single-cell suspensions were immediately processed by the University of Michigan Advanced Genomics Core. Libraries were constructed using the Chromium Next GEM Single Cell 3’ Gene Expression Kit v3.1 and Chromium Controller (10x Genomics) according to the manufacturer’s protocol. Each library was prepared with a target of 5,000 cells and processed independently. The resulting 10x-barcoded libraries were sequenced on an Illumina NovaSeq 6000 using an S4 flow cell with paired-end sequencing. Libraries were sequenced to a target depth of 100,000 reads per cell.

### Raw sequencing-data processing and reference construction

A custom reference containing the human GRCh38 genome and VA1 genome was generated using the Cell Ranger mkref pipeline version 7.1.0 (10x Genomics). The human component was based on GRCh38 reference version 2020-A with filtered GENCODE v32 annotations. The custom VA1 annotation contained positive-sense features corresponding to *nsp1ab* (nucleotides 39–4,222), *nsp1a* (nucleotides 39–2,699), and *capsid* (nucleotides 4,212–6,488) (NCBI’s VA1 Reference Sequence: NC_013060.1). Raw sequencing reads from each library were processed using the Cell Ranger count pipeline version 7.1.0 with intronic reads included and aligned to the combined GRCh38–VA1 reference. The resulting feature–barcode count matrices were used for downstream analysis in Seurat. Counts assigned to the VA1 *nsp1ab* and *capsid* features were used for analyses of viral transcript detection.

### Single-cell RNA sequencing transcriptomic analysis

A combined single-cell expression object containing mock-infected and VA1-infected libraries collected at 1 and 3 dpi was imported into R version 4.3.1 and converted to a Seurat object using readH5AD from the zellkonverter package version 1.10.1 and as.Seurat from Seurat version 5.2.1. Sample identity, inoculation condition, and collection time were retained in the object metadata. Cells were retained if they contained more than 2,000 detected features, fewer than 100,000 total counts, and less than 30% mitochondrial counts. Cells failing any one of these criteria were excluded from downstream analyses. Quality-control metrics were evaluated for each library before and after filtering. After filtering, the expression data were normalized using NormalizeData with the LogNormalize method and a scale factor of 10,000. Then, the 2,000 most variable features were identified using FindVariableFeatures with the vst selection method. Expression values for all genes were then centered and scaled using ScaleData. Principal-component analysis was then performed using RunPCA on the 2,000 variable features.

To prevent infection-induced transcriptional responses from influencing the identification of epithelial cell populations, clustering was performed using cells from the combined mock-infected libraries of both timepoints (7,335 total cells, 1,111 from 1 dpi and 6,224 from 3 dpi). A shared nearest-neighbor graph was generated using FindNeighbors with principal components 1–30. Clusters were identified using FindClusters at a resolution of 0.3. Uniform Manifold Approximation and Projection was performed using RunUMAP with principal components 1-30. Cluster markers were identified among the 2,000 variable features using FindAllMarkers with the Wilcoxon rank-sum test, a minimum detection fraction of 0.01, and an initial log2 fold-change threshold of 0.1. Markers with an average log2 fold change >1 and an adjusted p value <0.05 were retained for cell-type annotation. Cluster identities were then assigned by comparing marker expression with published fetal small-intestinal epithelial cell-type markers and expression profiles from the Human Protein Atlas (33). The initial analysis identified nine clusters. For downstream analyses, clusters 1 and 6 were consolidated as enterocytes; clusters 0, 2, 3, 5, and 7 were consolidated as progenitor cells; cluster 4 was annotated as goblet cells; and cluster 8 was annotated as enteroendocrine cells.

Cell-type annotations were transferred from the clustered mock reference to the VA1-infected query dataset containing both timepoints (5,408 total cells, 1,723 for 1 dpi and 3,685 for 3 dpi) using Seurat’s anchor-based label-transfer workflow. Transfer anchors were identified using FindTransferAnchors, with all features present in the mock reference supplied for anchor detection and all other parameters set to their Seurat defaults. Predicted cell-type labels were assigned using TransferData, with the manually curated annotations in the mock reference used as the reference labels. The resulting predicted.id assignments were added to the VA1 object metadata and used as the cell type identities. The annotated mock and VA1 objects were subsequently merged for downstream comparison.

VA1-feature detection was determined using raw unique molecular identifier counts assigned to the annotated VA1 nsp1ab and capsid features. Cells containing at least one viral transcript were considered VA1 transcript-positive and were classified as capsid-positive/nsp1ab-negative, nsp1ab-positive/capsid-negative, positive for both, or negative for both. Differential gene-expression analysis was performed using Seurat’s FindMarkers function with the Wilcoxon rank-sum test on normalized expression values, a minimum detection fraction of 0.01, and an initial log2 fold-change threshold of 0.1. Within each cell type and time point, VA1 transcript-positive cells from VA1-infected organoids were specified as ident.1 and compared with the corresponding cells from mock-infected organoids, specified as ident.2. Thus, positive average log2 fold-change values indicated higher expression in VA1 transcript-positive cells, whereas negative values indicated lower expression. Differentially expressed genes were retained if they had an absolute average log2 fold change >1 and an adjusted p value <0.05. Adjusted p values were calculated using Bonferroni correction across all features in the dataset. Viral transcripts were excluded from reported DEG lists. Shared DEGs were defined as genes that met these criteria in both enterocytes and progenitor cells at the same timepoint. Cell-type-exclusive DEGs were defined as genes meeting the DEG criteria in only one cell type. Shared and cell-type-exclusive DEG sets were identified using the R functions intersect and setdiff, respectively.

### Cell type proportion analysis

Cell-type proportions were calculated separately for mock-infected and VA1-infected samples at 1 and 3 dpi using the final four-population annotations: enterocytes, progenitor cells, goblet cells, and enteroendocrine cells. For each condition and timepoint, the proportion of each population was calculated as the number of cells assigned to that population divided by the total number of cells. Overall cell type distributions between mock- and VA1-infected samples were compared using two-sided Fisher’s exact tests applied to cell type-by-condition contingency tables. Individual cell type proportions were compared using two-sided two-sample tests for equality of proportions with continuity correction. Statistical significance was defined as p < 0.05.

### Gene Ontology Biological Processes & Transcription Factor Enrichment Analysis

Upregulated and downregulated DEG lists were analyzed separately at each time point for shared, enterocyte-exclusive, and progenitor-exclusive DEG sets. For Gene Ontology (GO) enrichment analysis, human gene symbols were converted to Entrez Gene identifiers using mapIds from the AnnotationDbi package and the org.Hs.eg.db annotation database, with keytype = “SYMBOL”, column = “ENTREZID”, and multiVals = “first”. Genes without a mapped Entrez identifier were excluded. GO Biological Process overrepresentation analysis was performed using the enrichGO function in clusterProfiler version 4.8.3, with OrgDb = org.Hs.eg.db, ont = “BP”, Bonferroni correction for multiple testing, and qvalueCutoff = 0.05. Because no custom universe was specified, the default enrichGO background was used. Redundant GO terms were reduced using clusterProfiler::simplify with a semantic-similarity cutoff of 0.7, with terms compared by adjusted p value and the term with the lowest adjusted p value retained within each group of semantically similar terms. Further redundant GO terms were excluded if they were a less significant term with at least 50% of its contributing DEGs overlapped with those of a more significant term. Entrez Gene identifiers associated with enriched terms were converted back to gene symbols for reporting. Terms with an adjusted p value < 0.05 were considered statistically significant, and the 5 most significant nonredundant terms were plotted.

Transcription factor enrichment analysis was performed by submitting the corresponding DEG lists to Enrichr (https://maayanlab.cloud/Enrichr/). Enrichment was assessed using the ENCODE and ChEA Consensus Transcription Factors from ChIP-X library. Enrichr-adjusted p values < 0.05 were considered statistically significant. Significant transcription-factor terms were ranked by adjusted p value and the 5 most significant terms were plotted.

### VA1 infection under lipid-depleted conditions

T84 and Caco-2 cells were seeded in 24-well plates at a density of 5 × 10^5^ cells/well. Two days after seeding, the standard growth medium was replaced with lipid-depleted medium prepared by substituting standard fetal bovine serum (FBS) with 10% lipid-depleted FBS (Alkali Scientific FB110). Cells were maintained in lipid-depleted medium for 24 hours before inoculation. The medium was then removed, and cells were inoculated with VA1 at a multiplicity of infection (MOI) of 100 genome copies per cell for 1 hour at 4°C. Next, cells were washed twice with PBS and replenished with lipid-depleted medium. Plates designated for the 0 dpi timepoint were immediately stored at −80 °C after washing. Remaining cultures were incubated at 37 °C with 5% CO₂ and collected at 1 dpi for T84 cells and 2 dpi for Caco-2 cells because of the slower VA1 growth kinetics in the latter. At the indicated time points, plates were stored at −80 °C until RNA extraction and VA1 genome copy quantification.

### Simvastatin treatment of T84 and Caco-2 cells

For T84 experiments, cells were seeded at 1.5 × 10^5^ cells per Transwell insert (Corning 3470) and cultured for 4 days to allow for polarization. Culture medium was replaced every other day. Cells were then treated apically with 100 µM simvastatin sodium salt (SVA, Cayman Chemical 10010345) prepared in PBS and diluted to the final concentration in lipid-depleted medium. Control cells were treated with an equivalent volume of PBS. The apical and basolateral compartments contained 200 μL and 500 μL of medium, respectively. Following a 24-hour pretreatment, cells were inoculated apically with VA1 at a multiplicity of infection (MOI) of 1,000 genome copies per cell. After adsorption for 1 hour at 37°C, the inoculum was removed, and cells were washed three times with PBS. Cells assigned to the 0 dpi time point were immediately lysed in Zymo Research TriReagent (R2050-1). For the 1 dpi timepoint, cultures were replenished with lipid-depleted medium containing 100 µM simvastatin hydroxy acid or the corresponding PBS vehicle control. At 1 dpi, the medium was removed, cells were lysed in Zymo Research TriReagent, and lysates were stored at −80 °C until RNA extraction and VA1 genome-copy quantification.

For Caco-2 experiments, cells were seeded in 24-well plates at a density of 5 × 10^5^ cells/well. Two days after seeding, standard growth medium was replaced with lipid-depleted medium containing 100 µM simvastatin lactone (Cayman Chemical 10010344) prepared as a stock solution in dimethyl sulfoxide (DMSO) and diluted in lipid-depleted medium. Control cells received the same final concentration of DMSO as the highest-dose treatment group. Following a 24-hour pretreatment, cells were inoculated with VA1 at an MOI of 100 genome copies per cell for 1 hour at 4 °C. The inoculum was removed, and cells were washed twice with PBS. Cells designated for the 0 dpi timepoint were immediately frozen at −80 °C. Remaining cultures were replenished with lipid-depleted medium containing the corresponding simvastatin concentration or DMSO vehicle control. At 2 dpi, the cells were frozen at −80 °C until RNA extraction and VA1 genome-copy quantification.

### U18cccA treatment and VA1 infection

T84 cells were seeded in 24-well plates at a density of 5 × 10^5^ cells/well. Two days after seeding, the standard growth medium was replaced with lipid-depleted medium. Cells were maintained in lipid-depleted medium for 24 hours before VA1 inoculation. Cells were then inoculated with VA1 at a MOI of 100 genome copies per cell for 1 hour at 4 °C. The inoculum was then removed, and cells were washed twice with PBS. Cells were subsequently maintained in lipid-depleted medium containing 10, 17.5, or 25 µM U18666A (Cayman Chemical 10009085). U18666A stock solutions were prepared in DMSO, and vehicle-control cells received the same final concentration of DMSO as the highest-dose treatment group. At 1 dpi, cells were frozen at −80 °C until RNA extraction and VA1 genome-copy quantification.

### 25-hydroxycholesterol treatment and VA1 infection

T84 cells and Caco-2 cells were seeded in 24-well plates at a density of 5 × 10^5^ cells/well. Three days after seeding, cells were then inoculated with VA1 at a MOI of 100 genome copies per cell for 1 hour at 4 °C. The inoculum was then removed, and cells were washed twice with PBS. Cells were subsequently maintained in medium containing 10 or 100 µM 25-hydroxycholesterol (25-HC) (Sigma-Aldrich H1015). 25-HC stock solutions were prepared in ethanol, and vehicle-control cells received the same final concentration of DMSO as the highest-dose treatment group. At 1 dpi for T84 cells and 2 dpi for Caco-2 cells, cells were frozen at −80 °C until RNA extraction and VA1 genome-copy quantification.

### RNA extraction and VA1 genome-copy quantification

RNA was extracted using the Direct-zol RNA Miniprep Plus kit (Zymo Research, R2072) according to the manufacturer’s instructions. VA1 genome copies were quantified by one-step quantitative reverse transcription PCR (RT-qPCR), as previously described (90). The following primer and probe sequences were used: VA1 forward primer, 5′-CTAGTGGTGGGGAAGAAC-3′; VA1 reverse primer, 5′-CCTTGGCTATTTGCTTTGC-3′; and VA1 probe, 5′-FAM-CCATGACTT-ZEN-TGCTTTGGACCTCCC-Iowa Black FQ-3′.

### Cell-viability measurement

Cell viability was assessed using WST-1 Cell Proliferation Reagent (Sigma-Aldrich, 5015944001). T84 or Caco-2 cells were seeded in 96-well plates at 9.6 × 10^4^ cells per well. Two days after seeding, the standard growth medium was replaced with lipid-depleted medium containing the indicated pharmacological treatment or corresponding vehicle control. Cells were incubated at 37 °C with 5% CO₂ for 48 hours, and the treatment was replenished after 24 hours. Cell viability was then measured according to the manufacturer’s instructions.

### Cellular cholesterol quantification

Cellular total cholesterol levels were measured using the Amplex Red Cholesterol Assay Kit (Invitrogen A12216). T84 and Caco-2 cells were seeded and treated with simvastatin or the corresponding vehicle control as described above. Cells were treated for 48 hours, with the treatment replenished after 24 hours. Cells were then detached using TrypLE Express Enzyme and washed three times with PBS. Cell pellets were resuspended in chloroform:methanol (2:1, v/v) and vortexed to facilitate lipid extraction. Following centrifugation, the organic phase was transferred to a new tube and evaporated under a stream of nitrogen. The dried lipid extracts were resuspended in the 1× reaction buffer supplied with the kit, and cholesterol-dependent fluorescence was measured according to the manufacturer’s instructions.

### Filipin staining of cells, imaging, and analysis

T84 cells were seeded in PhenoPlate^TM^-96 well plates (Revvity, 6055302). Three days post-seeding, cells were treated with 25 µM U18666A or an equivalent volume of DMSO (vehicle control) for 24 h. Cells were then rinsed once with PBS, fixed in 4% paraformaldehyde for 15 min at room temperature, and washed three times with PBS. Without permeabilization, cells were incubated with 50 µg/mL filipin III (Sigma-Aldrich, SAE0087) in PBS containing 10% FBS for 60 min at room temperature in the dark and washed three times with PBS. Images were acquired on a Yokogawa CQ1 spinning-disk confocal microscope with a 40×/0.95 NA objective. Filipin was excited at 405 nm and collected through a 447/60 nm filter. Each field was 0.162 µm per pixel, with 8 optical sections at 2.5 µm intervals. Four non-overlapping fields were imaged in each of 6 wells per condition. Images were corrected for dark current and flat-field, and coarse background was removed. For each field, the in-focus optical section with the most filipin structure was selected (maximum difference-of-Gaussians signal). Puncta were enhanced by Gaussian smoothing (σ = 1 pixel) followed by rolling-ball background subtraction (radius 1.6 µm). Candidate puncta were identified as regional maxima with a prominence of at least 25 and an intensity above 100 background-subtracted units. Filipin-labelled plasma membrane was excluded without cell segmentation by a radial test: candidates whose intensity stayed above 40% of their peak in two or more directions out to 1.6 µm were classified as membrane or cell junctions and rejected.

Puncta counts were normalized to tissue area, which was defined by a smoothed, thresholded filipin mask that excludes the lumen and gaps, and are reported as puncta per 1000 µm². Field values were averaged per well, and wells were the experimental unit (n = 6 per condition). Conditions were compared by two-sided Welch’s t-test and exact Mann– Whitney U test. Robustness was confirmed by repeating the analysis at intensity thresholds of 80 and 130. All parameters were fixed across images. Analysis was performed in Python 3.13 with NumPy, SciPy and scikit-image.

### Statistical analysis

For all experiments, data were analyzed either on R, Python 3.13 or Graphpad Prism 9 using the tests as indicated in methods and figure legends.

## Data availability statement

The raw and processed single-cell RNA-sequencing data generated in this study have been deposited in the NCBI Gene Expression Omnibus under accession GSE269641. The custom human astrovirus VA1 FASTA and GTF files used to construct the combined human–VA1 reference are provided with the GEO record.

## Acknowledgments

We thank all the members of the Wobus lab for regular discussions, Dr. Jonathan Golob (University of Michigan, MI, USA) for his help with initial pre-processing of the scRNAseq dataset, and Dr. Marcy Balunas (University of Michigan, MI, USA) for access to the nitrogen gas line during cholesterol assay sample processing.

This work was funded in part by NIH R21AI184493 and a University of Michigan pandemic relief grant to C.E.W. Research reported in this publication using the Yokogawa Cell Voyager 8000 instrumentation was supported by the Office of the Director, National Institutes of Health, under award number S10OD034245. A.R.R. was supported in part by NIH T32GM007544 and a University of Michigan’s Rackham Graduate Student Research Grant. M.M. was funded by NIH T32GM145304.

## References

1. Bosch A, Pintó RM, Guix S. 2014. Human Astroviruses. Clin Microbiol Rev 27:1048–1074.

2. De Benedictis P, Schultz-Cherry S, Burnham A, Cattoli G. 2011. Astrovirus infections in humans and animals – Molecular biology, genetic diversity, and interspecies transmissions. Infection, Genetics and Evolution 11:1529–1544.

3. GBD 2023 Diarrhoeal Disease and Enteric Infectious Diseases Collaborators. 2026. Global burden of enteric infectious diseases, diarrhoeal diseases, and corresponding aetiologies, 1990–2023: a systematic analysis for the Global Burden of Disease Study 2023. The Lancet Infectious Diseases 26:988–1009.

4. Prajapati B, Sikenis M, Singh AK, Diwan V, Mandal UK, Singh S, Pandey MK, Yadav K, Tiwari RR, Mishra PK, Yadav V, Nema RK. 2026. Prevalence of astrovirus-associated gastroenteritis among hospitalized pediatric patients in Asia: A systematic review and meta-analysis. Indian J Gastroenterol 45:1053–1067.

5. Fu J, Yu F, Li H, Shen L, Tian Y, Jia L, Zhang D, Yang P, Wang Q, Gao Z. 2023. Acute gastroenteritis outbreaks caused by human astrovirus, 1978-2021: A systematic review. Biosaf Health 5:120–125.

6. Koopmans MP, Bijen MH, Monroe SS, Vinjé J. 1998. Age-stratified seroprevalence of neutralizing antibodies to astrovirus types 1 to 7 in humans in The Netherlands. Clin Diagn Lab Immunol 5:33–37.

7. Kobayashi S, Kobayashi M, Araki K, Shinozaki T, Yanagawa Y. 1999. Antibody prevalence against astrovirus according to age groups. The Journal of the Japanese Association for Infectious Diseases 73:578–583.

8. Kurtz J, Lee T. 1978. Astrovirus gastroenteritis age distribution of antibody. Med Microbiol Immunol 166:227–230.

9. Wang J, Liu B, Liu S, Sun Y, He R, Lan Y, Liu L, Jiang H. 2025. Comparison of classic and novel human astrovirus MLB and VA seroprevalence in HIV and non-HIV cohorts in South China demonstrates high seroreactivity to classic human astrovirus, which is associated with HIV infection. Microbiol Spectr 13:e0083625.

10. Meyer L, Delgado-Cunningham K, Lorig-Roach N, Ford J, DuBois RM. 2021. Human Astrovirus 1-8 Seroprevalence Evaluation in a United States Adult Population. Viruses 13:979.

11. Mitchell DK, Matson DO, Cubitt WD, Jackson LJ, Willcocks MM, Pickering LK, Carter MJ. 1999. Prevalence of antibodies to astrovirus types 1 and 3 in children and adolescents in Norfolk, Virginia. Pediatr Infect Dis J 18:249–254.

12. Janowski AB, Owen MC, Dudley H, López T, Espinosa R, Elvin-Lewis M, Colichon A, Arias CF, Burbelo PD, Wang D. 2021. High Seropositivity Rate of Neutralizing Antibodies to Astrovirus VA1 in Human Populations. mSphere 6:e0048421.

13. Maximova OA, Weller ML, Krogmann T, Sturdevant DE, Ricklefs S, Virtaneva K, Martens C, Wollenberg K, Minai M, Moore IN, Sauter CS, Barker JN, Lipkin WI, Seilhean D, Nath A, Cohen JI. 2023. Pathogenesis and outcome of VA1 astrovirus infection in the human brain are defined by disruption of neural functions and imbalanced host immune responses. PLOS Pathogens 19:e1011544.

14. Holtz LR, Bauer IK, Jiang H, Belshe R, Freiden P, Schultz-Cherry SL, Wang D. 2014. Seroepidemiology of Astrovirus MLB1. Clin Vaccine Immunol 21:908–911.

15. Bengert B, Mehri S, Holliday M, Lennemann NJ. 2025. Endoplasmic reticulum-anchored nonstructural proteins drive human astrovirus replication organelle formation. PLoS Pathog 21:e1013538.

16. Noyvert D, Neves LX, Fominykh K, Hankinson J, Lindsey G, Lulla A, Emmott E, Lulla V. 2026. Viral protease-mediated polyprotein processing in human astroviruses. Npj Viruses 4:45.

17. Ali H, Noyvert D, Hankinson J, Lindsey G, Lulla A, Lulla V. 2024. The astrovirus N-terminal nonstructural protein anchors replication complexes to the perinuclear ER membranes. PLoS Pathog 20:e1011959.

18. Dong Q, Luo Y, Pan Y, Liu Y, Qin L, Ouyang K, Chen Y, Wei Z, Qin Y, Huang W. 2025. The C-terminal nsP1a protein of porcine astrovirus antagonizes interferon beta production by targets MAVS and IRF3 signaling. Vet Microbiol 305:110530.

19. Mehri S, Bengert B, Holliday M, Corliss L, Prevelige PE, Lennemann NJ. 2025. Identification of viral protease-dependent cleavage sites within the human astrovirus polyprotein. J Virol 99:e0132124.

20. Lentz A, López T, Ykema M, Tao YJ, DuBois RM, Arias CF. 2026. Human astroviruses: molecular virology. Microbiol Mol Biol Rev e0025525.

21. Lulla V, Firth AE. 2020. A hidden gene in astroviruses encodes a viroporin. Nat Commun 11:4070.

22. Noyvert D, Darr IM, Fominykh K, Hankinson J, Lukhovitskaya N, Firth AE, Lulla V. 2026. The dynamics and strategy of RNA replication in astroviruses. NAR Mol Med 3:ugag021.

23. Porto PS, Rivera A, Moonrinta R, Wobus CE. 2023. Chapter Two - Entry and egress of human astroviruses, p. 81–119. In Kielian, M, Roossinck, MJ (eds.), Advances in Virus Research. Academic Press.

24. Pedicino N, Kirkpatrick MG, Pablo MA, Gulman J, Heredia-Osuna N, Cortez V. 2026. Astroviruses use mucus secretion as a novel form of viral egress. PLoS Pathog 22:e1014683.

25. Kolawole AO, Wobus CE. 2020. Gastrointestinal organoid technology advances studies of enteric virus biology. PLoS Pathog 16:e1008212.

26. Adeniyi-Ipadeola G, Nwanosike H, Ramani S. 2023. Human intestinal organoids as models to study enteric bacteria and viruses. Curr Opin Microbiol 75:102362.

27. Gebert JT, Scribano F, Engevik KA, Perry JL, Hyser JM. 2023. Gastrointestinal organoids in the study of viral infections. Am J Physiol Gastrointest Liver Physiol 324:G51–G59.

28. Kolawole AO, Mirabelli C, Hill DR, Svoboda SA, Janowski AB, Passalacqua KD, Rodriguez BN, Dame MK, Freiden P, Berger RP, Vu D-L, Hosmillo M, O’Riordan MXD, Schultz-Cherry S, Guix S, Spence JR, Wang D, Wobus CE. 2019. Astrovirus replication in human intestinal enteroids reveals multi-cellular tropism and an intricate host innate immune landscape. PLoS Pathog 15:e1008057.

29. Triana S, Stanifer ML, Metz-Zumaran C, Shahraz M, Mukenhirn M, Kee C, Serger C, Koschny R, Ordoñez-Rueda D, Paulsen M, Benes V, Boulant S, Alexandrov T. 2021. Single-cell transcriptomics reveals immune response of intestinal cell types to viral infection. Mol Syst Biol 17:e9833.

30. Cieza RJ, Golob JL, Colacino JA, Wobus CE. 2021. Comparative Analysis of Public RNA-Sequencing Data from Human Intestinal Enteroid (HIEs) Infected with Enteric RNA Viruses Identifies Universal and Virus-Specific Epithelial Responses. Viruses 13:1059.

31. Platts-Mills JA, Babji S, Bodhidatta L, Gratz J, Haque R, Havt A, McCormick BJ, McGrath M, Olortegui MP, Samie A, Shakoor S, Mondal D, Lima IF, Hariraju D, Rayamajhi BB, Qureshi S, Kabir F, Yori PP, Mufamadi B, Amour C, Carreon JD, Richard S, Lang D, Bessong P, Mduma E, Ahmed T, Lima AA, Mason CJ, Zaidi AK, Bhutta Z, Kosek M, Guerrant RL, Gottlieb M, Miller M, Kang G, Houpt ER. 2015. Pathogen-specific burdens of community diarrhoea in developing countries (MAL-ED): a multisite birth cohort study. Lancet Glob Health 3:e564–e575.

32. Mirabelli C, Santos-Ferreira N, Gillilland MG, Cieza RJ, Colacino JA, Sexton JZ, Neyts J, Taube S, Rocha-Pereira J, Wobus CE. 2022. Human Norovirus Efficiently Replicates in Differentiated 3D-Human Intestinal Enteroids. J Virol 96:e0085522.

33. Elmentaite R, Ross ADB, Roberts K, James KR, Ortmann D, Gomes T, Nayak K, Tuck L, Pritchard S, Bayraktar OA, Heuschkel R, Vallier L, Teichmann SA, Zilbauer M. 2020. Single-Cell Sequencing of Developing Human Gut Reveals Transcriptional Links to Childhood Crohn’s Disease. Dev Cell 55:771–783.e5.

34. Stuart T, Butler A, Hoffman P, Hafemeister C, Papalexi E, Mauck WM, Hao Y, Stoeckius M, Smibert P, Satija R. 2019. Comprehensive Integration of Single-Cell Data. Cell 177:1888–1902.e21.

35. Cortez V, Boyd DF, Crawford JC, Sharp B, Livingston B, Rowe HM, Davis A, Alsallaq R, Robinson CG, Vogel P, Rosch JW, Margolis E, Thomas PG, Schultz-Cherry S. 2020. Astrovirus infects actively secreting goblet cells and alters the gut mucus barrier. Nat Commun 11:2097.

36. Ashburner M, Ball CA, Blake JA, Botstein D, Butler H, Cherry JM, Davis AP, Dolinski K, Dwight SS, Eppig JT, Harris MA, Hill DP, Issel-Tarver L, Kasarskis A, Lewis S, Matese JC, Richardson JE, Ringwald M, Rubin GM, Sherlock G. 2000. Gene Ontology: tool for the unification of biology. Nat Genet 25:25–29.

37. The Gene Ontology Consortium, Aleksander SA, Balhoff J, Carbon S, Cherry JM, Drabkin HJ, Ebert D, Feuermann M, Gaudet P, Harris NL, Hill DP, Lee R, Mi H, Moxon S, Mungall CJ, Muruganugan A, Mushayahama T, Sternberg PW, Thomas PD, Van Auken K, Ramsey J, Siegele DA, Chisholm RL, Fey P, Aspromonte MC, Nugnes MV, Quaglia F, Tosatto S, Giglio M, Nadendla S, Antonazzo G, Attrill H, dos Santos G, Marygold S, Strelets V, Tabone CJ, Thurmond J, Zhou P, Ahmed SH, Asanitthong P, Luna Buitrago D, Erdol MN, Gage MC, Ali Kadhum M, Li KYC, Long M, Michalak A, Pesala A, Pritazahra A, Saverimuttu SCC, Su R, Thurlow KE, Lovering RC, Logie C, Oliferenko S, Blake J, Christie K, Corbani L, Dolan ME, Drabkin HJ, Hill DP, Ni L, Sitnikov D, Smith C, Cuzick A, Seager J, Cooper L, Elser J, Jaiswal P, Gupta P, Jaiswal P, Naithani S, Lera-Ramirez M, Rutherford K, Wood V, De Pons JL, Dwinell MR, Hayman GT, Kaldunski ML, Kwitek AE, Laulederkind SJF, Tutaj MA, Vedi M, Wang S-J, D’Eustachio P, Aimo L, Axelsen K, Bridge A, Hyka-Nouspikel N, Morgat A, Aleksander SA, Cherry JM, Engel SR, Karra K, Miyasato SR, Nash RS, Skrzypek MS, Weng S, Wong ED, Bakker E, Berardini TZ, Reiser L, Auchincloss A, Axelsen K, Argoud-Puy G, Blatter M-C, Boutet E, Breuza L, Bridge A, Casals-Casas C, Coudert E, Estreicher A, Livia Famiglietti M, Feuermann M, Gos A, Gruaz-Gumowski N, Hulo C, Hyka-Nouspikel N, Jungo F, Le Mercier P, Lieberherr D, Masson P, Morgat A, Pedruzzi I, Pourcel L, Poux S, Rivoire C, Sundaram S, Bateman A, Bowler-Barnett E, Bye-A-Jee H, Denny P, Ignatchenko A, Ishtiaq R, Lock A, Lussi Y, Magrane M, Martin MJ, Orchard S, Raposo P, Speretta E, Tyagi N, Warner K, Zaru R, Diehl AD, Lee R, Chan J, Diamantakis S, Raciti D, Zarowiecki M, Fisher M, James-Zorn C, Ponferrada V, Zorn A, Ramachandran S, Ruzicka L, Westerfield M. 2023. The Gene Ontology knowledgebase in 2023. Genetics 224:iyad031.

38. Thomas PD, Ebert D, Muruganujan A, Mushayahama T, Albou L-P, Mi H. 2022. PANTHER: Making genome-scale phylogenetics accessible to all. Protein Science 31:8–22.

39. Kuleshov MV, Jones MR, Rouillard AD, Fernandez NF, Duan Q, Wang Z, Koplev S, Jenkins SL, Jagodnik KM, Lachmann A, McDermott MG, Monteiro CD, Gundersen GW, Ma’ayan A. 2016. Enrichr: a comprehensive gene set enrichment analysis web server 2016 update. Nucleic Acids Res 44:W90–97.

40. Chen EY, Tan CM, Kou Y, Duan Q, Wang Z, Meirelles GV, Clark NR, Ma’ayan A. 2013. Enrichr: interactive and collaborative HTML5 gene list enrichment analysis tool. BMC Bioinformatics 14:128.

41. Xie Z, Bailey A, Kuleshov MV, Clarke DJB, Evangelista JE, Jenkins SL, Lachmann A, Wojciechowicz ML, Kropiwnicki E, Jagodnik KM, Jeon M, Ma’ayan A. 2021. Gene Set Knowledge Discovery with Enrichr. Curr Protoc 1:e90.

42. Dunham I, Kundaje A, Aldred SF, Collins PJ, Davis CA, Doyle F, Epstein CB, Frietze S, Harrow J, Kaul R, Khatun J, Lajoie BR, Landt SG, Lee B-K, Pauli F, Rosenbloom KR, Sabo P, Safi A, Sanyal A, Shoresh N, Simon JM, Song L, Trinklein ND, Altshuler RC, Birney E, Brown JB, Cheng C, Djebali S, Dong X, Dunham I, Ernst J, Furey TS, Gerstein M, Giardine B, Greven M, Hardison RC, Harris RS, Herrero J, Hoffman MM, Iyer S, Kellis M, Khatun J, Kheradpour P, Kundaje A, Lassmann T, Li Q, Lin X, Marinov GK, Merkel A, Mortazavi A, Parker SCJ, Reddy TE, Rozowsky J, Schlesinger F, Thurman RE, Wang J, Ward LD, Whitfield TW, Wilder SP, Wu W, Xi HS, Yip KY, Zhuang J, Bernstein BE, Birney E, Dunham I, Green ED, Gunter C, Snyder M, Pazin MJ, Lowdon RF, Dillon LAL, Adams LB, Kelly CJ, Zhang J, Wexler JR, Green ED, Good PJ, Feingold EA, Bernstein BE, Birney E, Crawford GE, Dekker J, Elnitski L, Farnham PJ, Gerstein M, Giddings MC, Gingeras TR, Green ED, Guigó R, Hardison RC, Hubbard TJ, Kellis M, Kent WJ, Lieb JD, Margulies EH, Myers RM, Snyder M, Stamatoyannopoulos JA, Tenenbaum SA, Weng Z, White KP, Wold B, Khatun J, Yu Y, Wrobel J, Risk BA, Gunawardena HP, Kuiper HC, Maier CW, Xie L, Chen X, Giddings MC, Bernstein BE, Epstein CB, Shoresh N, Ernst J, Kheradpour P, Mikkelsen TS, Gillespie S, Goren A, Ram O, Zhang X, Wang L, Issner R, Coyne MJ, Durham T, Ku M, Truong T, Ward LD, Altshuler RC, Eaton ML, Kellis M, Djebali S, Davis CA, Merkel A, Dobin A, Lassmann T, Mortazavi A, Tanzer A, Lagarde J, Lin W, Schlesinger F, Xue C, Marinov GK, Khatun J, Williams BA, Zaleski C, Rozowsky J, Röder M, Kokocinski F, Abdelhamid RF, Alioto T, Antoshechkin I, Baer MT, Batut P, Bell I, Bell K, Chakrabortty S, Chen X, Chrast J, Curado J, Derrien T, Drenkow J, Dumais E, Dumais J, Duttagupta R, Fastuca M, Fejes-Toth K, Ferreira P, Foissac S, Fullwood MJ, Gao H, Gonzalez D, Gordon A, Gunawardena HP, Howald C, Jha S, Johnson R, Kapranov P, King B, Kingswood C, Li G, Luo OJ, Park E, Preall JB, Presaud K, Ribeca P, Risk BA, Robyr D, Ruan X, Sammeth M, Sandhu KS, Schaeffer L, See L-H, Shahab A, Skancke J, Suzuki AM, Takahashi H, Tilgner H, Trout D, Walters N, Wang H, Wrobel J, Yu Y, Hayashizaki Y, Harrow J, Gerstein M, Hubbard TJ, Reymond A, Antonarakis SE, Hannon GJ, Giddings MC, Ruan Y, Wold B, Carninci P, Guigó R, Gingeras TR, Rosenbloom KR, Sloan CA, Learned K, Malladi VS, Wong MC, Barber GP, Cline MS, Dreszer TR, Heitner SG, Karolchik D, Kent WJ, Kirkup VM, Meyer LR, Long JC, Maddren M, Raney BJ, Furey TS, Song L, Grasfeder LL, Giresi PG, Lee B-K, Battenhouse A, Sheffield NC, Simon JM, Showers KA, Safi A, London D, Bhinge AA, Shestak C, Schaner MR, Ki Kim S, Zhang ZZ, Mieczkowski PA, Mieczkowska JO, Liu Z, McDaniell RM, Ni Y, Rashid NU, Kim MJ, Adar S, Zhang Z, Wang T, Winter D, Keefe D, Birney E, Iyer VR, Lieb JD, Crawford GE, Li G, Sandhu KS, Zheng M, Wang P, Luo OJ, Shahab A, Fullwood MJ, Ruan X, Ruan Y, Myers RM, Pauli F, Williams BA, Gertz J, Marinov GK, Reddy TE, Vielmetter J, Partridge E, Trout D, Varley KE, Gasper C, The ENCODE Project Consortium, Overall coordination (data analysis coordination), Data production leads (data production), Lead analysts (data analysis), Writing group, NHGRI project management (scientific management), Principal investigators (steering committee), Boise State University and University of North Carolina at Chapel Hill Proteomics groups (data production and analysis), Broad Institute Group (data production and analysis), Cold Spring Harbor U of G Center for Genomic Regulation, Barcelona, RIKEN, Sanger Institute, University of Lausanne, Genome Institute of Singapore group (data production and analysis), Data coordination center at UC Santa Cruz (production data coordination), Duke University E University of Texas, Austin, University of North Carolina-Chapel Hill group (data production and analysis), Genome Institute of Singapore group (data production and analysis), HudsonAlpha Institute C UC Irvine, Stanford group (data production and analysis). 2012. An integrated encyclopedia of DNA elements in the human genome. Nature 489:57–74.

43. Lachmann A, Xu H, Krishnan J, Berger SI, Mazloom AR, Ma’ayan A. 2010. ChEA: transcription factor regulation inferred from integrating genome-wide ChIP-X experiments. Bioinformatics 26:2438–2444.

44. Yu G, Wang L-G, Han Y, He Q-Y. 2012. clusterProfiler: an R package for comparing biological themes among gene clusters. OMICS 16:284–287.

45. Wu T, Hu E, Xu S, Chen M, Guo P, Dai Z, Feng T, Zhou L, Tang W, Zhan L, Fu X, Liu S, Bo X, Yu G. 2021. clusterProfiler 4.0: A universal enrichment tool for interpreting omics data. Innovation 2.

46. Taniguchi T, Ogasawara K, Takaoka A, Tanaka N. 2001. IRF Family of Transcription Factors as Regulators of Host Defense. Annual Review of Immunology 19:623–655.

47. Dimitrov V, White JH. 2016. Species-specific regulation of innate immunity by vitamin D signaling. J Steroid Biochem Mol Biol 164:246–253.

48. Dannenberg J-H, David G, Zhong S, van der Torre J, Wong WH, Depinho RA. 2005. mSin3A corepressor regulates diverse transcriptional networks governing normal and neoplastic growth and survival. Genes Dev 19:1581–1595.

49. Savage KI, Gorski JJ, Barros EM, Irwin GW, Manti L, Powell AJ, Pellagatti A, Lukashchuk N, McCance DJ, McCluggage WG, Schettino G, Salto-Tellez M, Boultwood J, Richard DJ, McDade SS, Harkin DP. 2014. Identification of a BRCA1-mRNA splicing complex required for efficient DNA repair and maintenance of genomic stability. Mol Cell 54:445–459.

50. Balamurugan K, Sterneck E. 2013. The Many Faces of C/EBPδ and their Relevance for Inflammation and Cancer. International Journal of Biological Sciences 9:917–933.

51. Litovchick L, Sadasivam S, Florens L, Zhu X, Swanson SK, Velmurugan S, Chen R, Washburn MP, Liu XS, DeCaprio JA. 2007. Evolutionarily Conserved Multisubunit RBL2/p130 and E2F4 Protein Complex Represses Human Cell Cycle-Dependent Genes in Quiescence. Molecular Cell 26:539–551.

52. Wierstra I, Alves J. 2007. FOXM1, a typical proliferation-associated transcription factor 388:1257–1274.

53. Zhang J, Tam W-L, Tong GQ, Wu Q, Chan H-Y, Soh B-S, Lou Y, Yang J, Ma Y, Chai L, Ng H-H, Lufkin T, Robson P, Lim B. 2006. Sall4 modulates embryonic stem cell pluripotency and early embryonic development by the transcriptional regulation of Pou5f1. Nat Cell Biol 8:1114–1123.

54. Noah R, Pan Q, Ruiz AR, Jorge DMDM, Raya-Sandino A, Carreteiro de Oliveira MS, Hafner A, McConnachie M, Sexton JZ, Nusrat A, Wobus CE. 0. Non-classical human astrovirus VA1 induces cell line-specific increases in intestinal epithelial permeability associated with claudin remodeling. J Virol 0:e00845–26

55. Ward NC, Watts GF, Eckel RH. 2019. Statin Toxicity. Circ Res 124:328–350.

56. Lu F, Liang Q, Abi-Mosleh L, Das A, De Brabander JK, Goldstein JL, Brown MS. Identification of NPC1 as the target of U18666A, an inhibitor of lysosomal cholesterol export and Ebola infection. eLife 4:e12177.

57. Zhang J, Zhu Y, Wang X, Wang J. 2023. 25-hydroxycholesterol: an integrator of antiviral ability and signaling. Front Immunol 14:1268104.

58. Liang Z, Xing S, Chen Y, Liu M, Chang S, Cai Y, Xiong Q, Xu Y, Liu J. 2026. Dynamic regulatory mechanism of cholesterol 25-hydroxylase-mediated antiviral response in RNA virus infections. Front Cell Infect Microbiol 16:1790475.

59. Costantini V, Mboko WP, Wall H, Huynh K, Montmayeur AM, Park GW, Xu R, Vinjé J, Chhabra P. 2025. 25-Hydroxycholesterol restricts human norovirus replication in human intestinal enteroids. J Virol 99:e0110925.

60. Shawli GT, Adeyemi OO, Stonehouse NJ, Herod MR. 2019. The Oxysterol 25-Hydroxycholesterol Inhibits Replication of Murine Norovirus. Viruses 11:97.

61. Huang H-I, Lin J-Y, Chiang H-C, Huang P-N, Lin Q-D, Shih S-R. 2020. Exosomes Facilitate Transmission of Enterovirus A71 From Human Intestinal Epithelial Cells. J Infect Dis 222:456–469.

62. Haimovich G, Dasgupta S, Gerst JE. 2021. RNA transfer through tunneling nanotubes. Biochem Soc Trans 49:145–160.

63. Matsushima Y, Durkee-Shock J, Alves DA, Nagata BM, Kim SA, Chan EC, Chaimongkol N, Ishiyama R, Barton K, Han A, Kelly LA, Ogbogu PU, Kumar S, Xu X, Briggs KC, Murakami K, Cohen JI, Keller MD, Sosnovtsev SV, Green KY. 2026. Single-cell profiling of the duodenum during chronic human norovirus infection. Science Advances 12:eaej2676.

64. Bomidi C, Robertson M, Coarfa C, Estes MK, Blutt SE. 2021. Single-cell sequencing of rotavirus-infected intestinal epithelium reveals cell-type specific epithelial repair and tuft cell infection. Proc Natl Acad Sci U S A 118:e2112814118.

65. Triana S, Metz-Zumaran C, Ramirez C, Kee C, Doldan P, Shahraz M, Schraivogel D, Gschwind AR, Sharma AK, Steinmetz LM, Herrmann C, Alexandrov T, Boulant S, Stanifer ML. 2021. Single-cell analyses reveal SARS-CoV-2 interference with intrinsic immune response in the human gut. Mol Syst Biol 17:e10232.

66. Colgan SP, Furuta GT, Taylor CT. 2020. Hypoxia and Innate Immunity: Keeping Up with the HIFsters. Annu Rev Immunol 38:341–363.

67. Jacobs SO, Muenchau S, Uckeley ZM, Passarelli GV, David A, Briggs S, Burke JM, Stanifer ML, Boulant S. 2025. Hypoxia increases susceptibility of human intestinal epithelial cells to rotavirus infection through repression of interferon induction. Gut Microbes 17:2560593.

68. Lee BR, Paing MH, Sharma-Walia N. 2021. Cyclopentenone Prostaglandins: Biologically Active Lipid Mediators Targeting Inflammation. Front Physiol 12:640374.

69. Boisvert M, Côté S, Vargas A, Pasvanis S, Bounou S, Barbeau B, Dumais N. 2008. PGJ2 antagonizes NF-kappaB-induced HIV-1 LTR activation in colonic epithelial cells. Virology 380:1–11.

70. Lee H, Ko G. 2016. Antiviral effect of vitamin A on norovirus infection via modulation of the gut microbiome. Sci Rep 6:25835.

71. Lai X, Wu A, Yu B, Yan H, Luo J, Zheng P, Yu J, Chen D. 2024. Retinoic acid alleviates rotavirus-induced intestinal damage by regulating redox homeostasis and autophagic flux in piglets. Anim Nutr 16:409–421.

72. Kim Y, Chang K-O. 2011. Inhibitory effects of bile acids and synthetic farnesoid X receptor agonists on rotavirus replication. J Virol 85:12570–12577.

73. He H, Shen H, Zhang C, Zhang K, Qu Y, Nie J, Jia A, Zhang J, Liu Z. 2026. Lithocholic acid exerts antiviral activity against porcine epidemic diarrhea virus by enhancing TGR5-mediated type III interferon production. Microb Pathog 210:108187.

74. Makimaa H, Ingle H, Wang L, Aggarwal S, Deng H, Kennedy EA, Foster L, Li Y, Baldridge MT. 2026. MDA5-MAVS and interferon-lambda signaling in the intestinal epithelium limit murine astrovirus infection. Mucosal Immunol 19:1749–1758.

75. Yan H, Ye Y, Zhao H, Zuo H, Li Y. 2022. Single-Cell RNA Sequencing for Analyzing the Intestinal Tract in Healthy and Diseased Individuals. Front Cell Dev Biol 10.

76. Kalugotla G, Marmerstein V, Baldridge MT. 2024. Regulation of host/pathogen interactions in the gastrointestinal tract by type I and III interferons. Curr Opin Immunol 87:102425.

77. Ma LJ, Acero LF, Zal T, Schluns KS. 2009. Trans-presentation of IL-15 by intestinal epithelial cells drives development of CD8αα IELs. J Immunol 183:1044–1054.

78. Di Sabatino A, Ciccocioppo R, Cupelli F, Cinque B, Millimaggi D, Clarkson MM, Paulli M, Cifone MG, Corazza GR. 2006. Epithelium derived interleukin 15 regulates intraepithelial lymphocyte Th1 cytokine production, cytotoxicity, and survival in coeliac disease. Gut 55:469–477.

79. Reinecker HC, MacDermott RP, Mirau S, Dignass A, Podolsky DK. 1996. Intestinal epithelial cells both express and respond to interleukin 15. Gastroenterology 111:1706–1713.

80. Tamzalit F, Barbieux I, Plet A, Heim J, Nedellec S, Morisseau S, Jacques Y, Mortier E. 2014. IL-15.IL-15Rα complex shedding following trans-presentation is essential for the survival of IL-15 responding NK and T cells. Proc Natl Acad Sci U S A 111:8565–8570.

81. Wu X, Dao Thi VL, Huang Y, Billerbeck E, Saha D, Hoffmann H-H, Wang Y, Silva LAV, Sarbanes S, Sun T, Andrus L, Yu Y, Quirk C, Li M, MacDonald MR, Schneider WM, An X, Rosenberg BR, Rice CM. 2018. Intrinsic Immunity Shapes Viral Resistance of Stem Cells. Cell 172:423–438.e25.

82. Jiang L, Tang A, Song L, Tong Y, Fan H. 2023. Advances in the development of antivirals for rotavirus infection. Front Immunol 14:1041149.

83. Cuadras MA, Greenberg HB. 2003. Rotavirus infectious particles use lipid rafts during replication for transport to the cell surface in vitro and in vivo. Virology 313:308–321.

84. Ayyar BV, Apostol CV, Dave JJ, Kaundal S, Kendra JA, Neill FH, Ettayebi K, Maher S, Anish R, Parra GI, Larson G, Atmar RL, Crawford SE, Prasad BVV, Estes MK. 2025. Functional diversity in GII.4 norovirus entry: HBGA binding and capsid clustering dynamics. Proc Natl Acad Sci U S A 122:e2517493122.

85. Stoeck IK, Lee J-Y, Tabata K, Romero-Brey I, Paul D, Schult P, Lohmann V, Kaderali L, Bartenschlager R. 2017. Hepatitis C Virus Replication Depends on Endosomal Cholesterol Homeostasis. J Virol 92:e01196–17.

86. Bub T, Hargest V, Tan S, Smith M, Vazquez-Pagan A, Flerlage T, Brigleb P, Meliopoulos V, Lindenbach B, Ramanathan HN, Cortez V, Crawford JC, Schultz-Cherry S. 2023. Astrovirus replication is dependent on induction of double-membrane vesicles through a PI3K-dependent, LC3-independent pathway. J Virol 97:e0102523.

87. Bustamante-Madrid P, Barbáchano A, Albandea-Rodríguez D, Rodríguez-Cobos J, Rodríguez-Salas N, Prieto I, Burgos A, Martínez de Villarreal J, Real FX, González-Sancho JM, Larriba MJ, Lafarga M, Muñoz A, Fernández-Barral A. 2024. Vitamin D opposes multilineage cell differentiation induced by Notch inhibition and BMP4 pathway activation in human colon organoids. Cell Death Dis 15:301.

88. Basak O, Beumer J, Wiebrands K, Seno H, van Oudenaarden A, Clevers H. 2017. Induced Quiescence of Lgr5+ Stem Cells in Intestinal Organoids Enables Differentiation of Hormone-Producing Enteroendocrine Cells. Cell Stem Cell 20:177–190.e4.

89. Adeniyi-Ipadeola GO, Hankins JD, Kambal A, Zeng X-L, Patil K, Poplaski V, Bomidi C, Nguyen-Phuc H, Grimm SL, Coarfa C, Stossi F, Crawford SE, Blutt SE, Speer AL, Estes MK, Ramani S. 2024. Infant and adult human intestinal enteroids are morphologically and functionally distinct. mBio 15:e0131624.

90. Janowski AB, Bauer IK, Holtz LR, Wang D. 2017. Propagation of Astrovirus VA1, a Neurotropic Human Astrovirus, in Cell Culture. J Virol 91:e00740–17.

